# Standing genetic variation drives polygenic adaptation to different environmental shifts

**DOI:** 10.64898/2026.09.14.751342

**Authors:** Yuna Zhang, Markus G Stetter

**Affiliations:** Institute for Plant Sciences, University of Cologne, Cologne, Germany

**Keywords:** Polygenic adaptation, standing genetic variation, environmental change, environmental fluctuation, allele frequency

## Abstract

Most organisms are well adapted to the environment they have been exposed to for many generations. However, changing environments pose existential threats to populations, unless they are able to adapt. Hence, a central challenge in evolutionary genetics is to understand the adaptation to complex environmental changes. As most traits are controlled by a large number of loci with varying effects on a trait, it is challenging to understand the explicit role of genetic changes during such polygenic adaptation. We use forward-in-time simulations to investigate the evolutionary dynamics of phenotypic and genetic adaptation in populations facing different environmental shifts. Specifically, we simulated sudden, gradual, and fluctuating environmental shifts for multiple trait architectures. Our results show distinct evolutionary paths across different types of environmental shifts and a higher extinction risk during sudden and fluctuating environmental shifts than during gradual shifts. Comparing the contribution of mutations from different sources highlights the critical role of standing genetic variation in driving phenotypic adaptation. We summarize allele frequency trajectories by clustering them by their temporal pattern and reveal how large- and small-effect mutations jointly shape the successful adaptation to changing environments. Additionally, we trained a convolutional neural network (CNN) on ‘genetic architecture matrices’ of populations to jointly infer the type and magnitude of environmental changes and the mutational effect size distribution. The CNN was able to predict all three input parameters with very high accuracy, even on unseen parameter combinations. Our results demonstrate the impact of ecological change on the evolutionary outcome and the assorted mutational changes that enable successful adaptation.

## Introduction

Environmental change is pervasive and accelerating in the face of climate change (Walther *et al*. 2002; Hoffmann and Sgrò 2011). Populations inhabiting altered environments must either adapt to the novel environments or face extinction. Understanding whether populations can adapt to a changing environment and how, is a central question in evolutionary biology and of major relevance for species conservation (Hancock *et al*. 2025). However, populations experience diverse types of environmental shifts, including sudden environmental shifts (e.g., extreme climatic events (Baeckens and Donihue 2025) or sudden exposure to antibiotics (Windels *et al*. 2020)), gradual shifts (e.g., global warming (Radchuk *et al*. 2019)), and temporal fluctuations (e.g., seasonal shifts (Behrman *et al*. 2015)). Because these shifts differ in the tempo and direction of selection, they impose distinct selective pressures and are expected to shape both the trajectory and outcome of adaptation differently. Critically, the pace of environmental change has intensified markedly over the past two centuries, through industrialization and massive population growth. For short-lived organisms, such as annual plants, this represents hundreds of generations of environmental shifts (Kinnison and Hairston 2007; Tellier *et al*. 2024). Despite awareness of rapid environmental change, it is predicted that directional environmental change will continue for a long time (Grant *et al*. 2025). Therefore, it is important to understand if and how populations can adapt to continued rapid environmental change.

Adaptation is the evolutionary process through which populations track the optimal phenotype that best fits the environment in response to selection (Lande 1976; Futuyma 2005). In natural systems, this process typically manifests as phenotypic shifts in traits with continuous variation, as observed for beak size in Darwin’s finches (Grant and Grant 2002), or flowering time in plants(Anderson *et al*. 2012). Although population genetics often emphasizes adaptation driven by a few large-effect mutations (Smith and Haigh 1974; Orr 1998), the adaptation of such quantitative traits is mostly polygenic (Pritchard *et al*. 2010; Pritchard and Di Rienzo 2010). This perspective is reinforced by extensive genome-wide association studies, which have revealed that the majority of traits are governed by a large number of loci rather than a few (Boyle *et al*. 2017; Buniello *et al*. 2019; Tautz *et al*. 2026). Consequently, the study of quantitative traits provides an essential framework for understanding the dynamics of polygenic adaptation, where coordinated frequency shifts across many loci move the population mean toward a new optimum (Barghi *et al*. 2020). However, the explicit roles and trajectories of adaptive genetic variants during this process remain poorly resolved.

Adaptation can occur through selection on *de novo* mutations, or on standing genetic variation present in the population before environmental change. Despite the role of new mutations in driving evolutionary change (Orr 2005), mounting empirical evidence indicates that standing genetic variation plays an important role during rapid adaptive responses (Lai *et al*. 2019; Bitter *et al*. 2019; Yang *et al*. 2025; Stetter *et al*. 2018). Mutations that arose before the environmental change are present at the onset of the directional selection pressure. Small-effect standing variants are often nearly neutral or weakly deleterious in the original environments, allowing them to reach intermediate frequencies. But these alleles might become beneficial following environmental changes, leading to rapid increases in frequency under positive selection (Hermisson and Pennings 2005; Barrett and Schluter 2008; Matuszewski *et al*. 2015). Unlike small-effect variants, large-effect mutations are typically at very low frequency due to strong purifying selection. Yet, pre-existing variation can still respond more rapidly to directional selection than large-effect *de novo* mutations, as waiting times for new beneficial mutations to arise can be long (Barrett and Schluter 2008). Even once one arises, its probability of fixation depends strongly on the genetic background into which it arises (Chevin and Hospital 2008). Hence, the balance between effect size and starting frequency is an important driver of adaptation (Hermisson and Pennings 2005). Individual standing variants often leave only soft selective sweeps or no sweep at all, depending on the allele age and frequency change during adaptation (Hermisson and Pennings 2005; Barrett and Schluter 2008). Often, the polygenic combination of small-effect mutations with small frequency shifts leaves a complicated pattern of population diversity across the genome rather than distinct signatures at individual loci (Pritchard *et al*. 2010). Despite its importance, the relative contribution of standing genetic variation in response to environmental change remains poorly understood.

The type of environmental change can have a substantial impact on the genetic makeup of a population. Sudden environmental shifts lead to an extreme change in the phenotypic optimum, often beyond the phenotypic variance, and require large genetic changes (Jain and Stephan 2015, 2017; Stetter *et al*. 2018; Hayward and Sella 2022). Gradual change can still reach equally strong shifts, but smaller shifts per generation alter the composition of genetic diversity and the observed allele combinations (Bürger and Lynch 1995; Kopp and Hermisson 2009a,b; Matuszewski *et al*. 2015; Pahujani *et al*. 2026). Experimental evolution in *Caenorhabditis elegans* showed that sudden environmental shifts rapidly increase allele frequency at large-effect standing genetic variants, whereas gradual environmental change initially favored small-effect variants (Guzella *et al*. 2018). Understanding the impact of the type of environmental change on the genetic outcome will help understand vulnerabilities of populations in the face of change.

In this study, we aim to elucidate the consequences of rapid environmental change. We employed individual-based forward-in-time simulations (Messer 2013) to investigate the potential of polygenic adaptation across multiple environmental shifts. We simulated 400 years of environmental change for an annual organism (many plants), representing a human-driven environmental change through urbanization and industrialization. Specifically, we compared adaptation under sudden, gradual, and fluctuating shifts with different magnitudes of change for different trait types and show that the type and strength of change strongly alter the adaptive success of a population. By integrating phenotypic and genetic responses across a broad parameter space, our study underlines the contribution of standing genetic variation and characteristic allele frequency trajectories during polygenic adaptation. We train a machine learning model and show that genetic data from a single time point after directional change is sufficient to reveal the adaptation history of the population.

## Methods

### Quantitative trait simulation

We simulated an outcrossing, diploid population with discrete generations (Wright-Fisher model). Specifically, we used a constant population size of *N* = 10^5^, similar to observed plant populations (Durvasula *et al*. 2017). We used realistic recombination rates varying across the chromosome (chromosome 4) by integrating empirical data from the model plant *Arabidopsis thaliana* (Salomé *et al*. 2012), yielding a chromosome-wide average of 3.23 × 10^−8^ per base pair per gamete (Figure S1). We used the estimated mutation rate of *A. thaliana* (*µ* = 7 × 10^−9^ per site per generation) (Ossowski *et al*. 2010) and the gene structure of chromosome 4 (∼ 18.6 Mb) from the reference genome (Swarbreck *et al*. 2008). With this, we assumed that 10% of mutations within genes and 1% of mutations in non-genic regions can affect a trait (Figure S1). Hence, the absolute mutation rate was 1492 trait-affecting mutations per generation.

We simulated quantitative traits with different effect sizes of new mutations that were exposed to different combinations of changing trait optima. The trait value *z* is defined as the additive sum of the effect sizes of all alleles present in an individual (De Vladar and Barton 2014). An individual’s fitness *w* is thus expressed as (Bürger 2000):

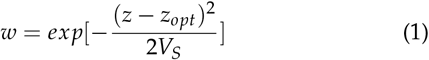

where *z*_*opt*_ is the trait optimum, and 1/*V*_*S*_ represents the strength of stabilizing selection.

Effect sizes of new mutations were drawn from a normal distribution, *N*(0, *ω*^2^), where *ω* denotes the standard deviation of the effect sizes of new mutations. As 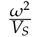 determines the effective selection strength acting on a trait (Pahujani *et al*. 2026), we fixed *V*_*S*_ = 10 and varied *ω* across six values (0.05, 0.1, 0.2, 0.3, 0.4, and 0.5) to simulate a range of trait architectures.

For all parameter combinations, we simulated a starting population with a quantitative trait subject to stabilizing selection (*z*_*opt*_ = 0) for a burn-in period of 10 N (10^6^) generations to create an equilibrium population in mutation-selection-drift equilibrium (Haller *et al*. 2019). We started recording data after the burn-in, recording 99 generations (starting at generation -99) of stabilizing selection, followed by different environmental change trajectories. To investigate how environmental trajectories affect adaptation, each starting population was subjected to three types of environmental shifts, namely a sudden, a gradual, and a fluctuating shift of the phenotypic optimum, beginning at generation 0 (Figure S2). To make the trajectories comparable between the types of environmental shifts, we kept the final magnitude of shift after 400 generations equal across shifts. In the ‘sudden shift’, the optimum was set instantaneously to the final optimum at generation 0 and remained constant thereafter. In the ‘gradual shift’, the optimum shifted linearly from 0 to the new optimum value over 400 generations (0-399). In the ‘fluctuating shift’, the optimum at each generation was sampled from a normal distribution centered on the same linear trajectory as the gradual shift, with a standard deviation of one. From generation 400 to 500, the optimum was fixed, inducing stabilizing selection in all three types of environmental shifts. This enabled the comparison of the long-term impact of differences due to past environmental change, even when the environment was constant. Across all three shifts, we studied eight shift magnitudes (1, 2, 3, 4, 5, 10, 20, 50) that represent strengths of environmental change from weak (1) to very strong (50). In addition, we simulated a static environment where the optimum was fixed at zero throughout the observed time period, representing an environment with continuous stabilizing selection.

In total, we simulated 150 parameter combinations: 6 different traits represented through the change in *ω* × 25 environmental conditions (3 ‘environmental shifts’ × 8 magnitudes of change + 1 static). We carried out 100 independent simulations for each combination of parameters using SLiM 4 (Haller and Messer 2023). For each of the 15,000 simulations, we recorded the phenotypic and genotypic data at every generation for a total of 600 generations.

### Simulations without standing variation or de novo mutations

To assess whether standing variation was necessary for adaptation rather than merely accounting for most of the phenotypic change, we ran two additional sets of simulations in which one source of genetic variation was removed. To remove standing variation, we initialized a population without any genetic variation, began the optimum shift in its first generation (indexed as generation 0), and produced mutations after the optimum shift. These populations evolved only using *de novo* mutations. To remove *de novo* mutations, we used the populations at generation 0 from the aforementioned simulations (the factual simulations with both standing and *de novo* variants) to retain their complete standing variation, and set the mutation rate to zero onward. These populations evolved only using standing mutations. We restricted this analysis to the gradual shift with magnitude = 50 and *ω* = 0.1, a magnitude at which *de novo* mutation has greater opportunity to contribute to the phenotypic change and the *ω* value with which populations successfully adapted by generation 400. Therefore, these counterfactual simulations constitute conservative tests. All other population parameters and the optimum trajectory were kept the same as in previous simulations (with standing and new mutations), with 100 replicates per design. We compared the dynamics of mean fitness from generation 0 to 500. Because removing one source alters the dynamics of the other, the phenotypic values under the two counterfactual simulations are not complementary.

### Analysis of evolutionary outcomes

We recorded three population-level statistics at every generation: mean phenotype, genetic variance, and mean fitness. Because individual phenotype is defined as the additive sum of allelic effects, genetic variance is the variance of individual phenotypes in a population. Mean phenotypic lag was calculated as the difference between the optimum and the mean phenotype at each generation 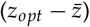.

We assessed the evolutionary outcome of each population based on mean fitness at generation 400, 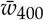: a population was considered successfully adapted if 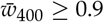; extinct if 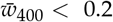; and surviving but not yet fully adapted if 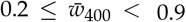. Populations that crossed the extinction threshold before generation 400 were assigned the 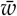 recorded at the generation they first crossed it for subsequent linear modeling.

To assess the impact of environmental shift, magnitude of the shift, and *ω* on the evolutionary outcomes, we fitted a linear model:

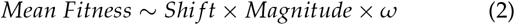

using permutation tests via the “lmp” function in the “lmPerm” R package, which does not require strict parametric assumptions. We partitioned variance among the model terms with a type-III ANOVA via the “Anova” function in the “car” package and estimated their marginal effects using partial *η*^2^ via the “eta_squared” function from the “effectsize” package.

### Phenotypic contribution of mutations

We quantified the phenotypic contribution of mutations as 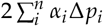, where *α*_*i*_ represents the allelic effect of the *i*-th mutation, Δ*p*_*i*_ denotes the change in allele frequency, and *n* is the number of mutations; the coefficient of 2 is for diploids. Using this statistic, we compared the contribution of standing genetic variants present at generation 0 and *de novo* mutations arising after generation 0 for each parameter combination. In addition, we calculated the phenotypic changes resulting from fixed mutations (*p* = 1 between generation 1-400) and lost mutations (*p* = 0 between generation 1-400).

### Clustering allele frequency trajectories

To study how standing variation drives adaptation, we assigned all standing alleles based on the similarity of their frequency trajectories. For each of the 150 parameter combinations, we compiled allele frequencies across all generations (0 to 500) from all 100 replicates, then centered each trajectory by subtracting the allele’s starting frequency at generation 0, so that clustering reflects the magnitude and direction of frequency change rather than absolute frequency. We performed Principal Component Analysis (PCA) (“prcomp”, “stats” R package) on these centered trajectories and retained the first three PCs, which collectively explained more than 90% of the total variance, for subsequent k-means clustering (“kmeans”, “stats” R package). The optimal number of clusters was set to *k* = 8, identified by computing the marginal reduction in total within-cluster sum of squares (*WSS*_*k* −1_ −*WSS*_*k*_) across *k* ∈ [2, 20] and selecting the point at which this reduction began to plateau (Figure S3).

To quantify whether the phenotypic contribution of standing genetic variation was dominated by a single cluster or broadly distributed across clusters, we calculated “the effective number of clusters”, which is the exponential of the Shannon entropy of the normalized cluster contributions. We first excluded cluster 8, because it assembles a wide range of allele frequency shifts, and normalized the contribution (*c*_*i*_) of the remaining seven clusters to obtain relative weights (summing to 1):

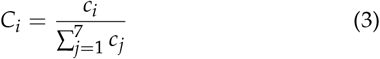

Then calculated the Shannon entropy (Shannon 1948):

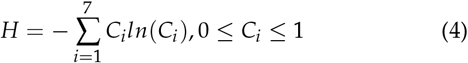

The Hill number at the order (*q*) of 1 is calculated as *exp*(*H*) (Jost 2006), which measures the number of equally contributing clusters that would yield the same distribution of contribution as observed.

**Table 1.** Overview of parameters and variables used in this study.

| Variable | Description |
| --- | --- |
| $N$ | Population size |
| $\mu$ | Mutation rate per site per generation |
| $w$ | Fitness |
| $V_S$ | Variance of the fitness function (strength of stabilizing selection) |
| $z$ | Phenotypic value |
| $z_{opt}$ | Optimal phenotypic value |
| $V_G$ | Genetic variance of a phenotype |
| $\omega$ | Standard deviation of effect size of new mutations |
| $\alpha$ | Effect size of alleles |
| $p$ | Allele frequency |

### Deep learning: multi-task inference via two-dimensional convolutional neural networks

For each simulated population, we summarized the genetic information at generation 400 as a ‘genetic architecture matrix’. This 20 × 40 matrix has rows corresponding to 20 equal-width allele frequency bins and columns corresponding to 40 equal-width effect-size bins symmetric around zero (including 20 negative bins and 20 positive bins, and the bin width is 1/20 of the largest absolute effect size segregating in that population). Effect-size bins were therefore defined relative to each population rather than on a common absolute scale, so that populations differing in *ω* could not be distinguished by the range of occupied columns alone. Each matrix element represents the proportion of segregating alleles at that frequency-effect coordinate, such that the matrix sums to 1 (see Figure 5A). The full dataset comprised 15,000 matrices, 4,800 for each of the three environmental shifts (100 replicates × 6 values of *ω* × 8 magnitudes) and 600 for the static environment (100 replicates × 6 values of *ω* × 1 magnitude (=0)).

We implemented a multi-task two-dimensional convolutional neural network (2D CNN) in Python using TensorFlow version 2.19 (Abadi *et al*. 2015) to jointly infer three quantities from each genetic architecture matrix: the type of environmental shift (classification task), the magnitude of environmental shift (regression task), and the standard deviation of the mutational effect size distribution, *ω* (regression task). The network consisted of convolutional layers for feature extraction, followed by shared fully connected layers and task-specific output heads.

We randomly sampled 10% of the 15,000 matrices as a heldout test set, stratified by environmental shift type so that each of the three non-static shift types contributed 480 matrices, and the static type contributed 60. The remaining 13,500 matrices formed the tuning set, on which we performed hyperparameter search, cross-validation, and final model fitting. The test set was evaluated once after the final model fitting.

We selected hyperparameters using a grid search over 32 combinations of learning rate (10^−4^, 5 × 10^−4^, 10^−3^, 10^−2^), dropout rate (0.2, 0.3, 0.4, 0.5), dense-layer sizes (256 and 128, or 128 and 64). For each combination of hyperparameters, we performed 10-fold cross-validation (Greener *et al*. 2022). We partitioned the tuning set into 10 subsets of 1,350 matrices, stratified by environmental shift type. In each fold, one subset served as the evaluation split, and the remaining nine were split into 10,935 matrices for training and 1,215 for early stopping, again stratified by shift type. We standardized the input matrices with a single global mean and standard deviation, and the magnitude and *ω* targets with their own means and standard deviations, all computed on the training split and applied to the other two splits to prevent data leakage. Predictions were transformed back to the original scale before scoring. Models were trained for up to 100 epochs by minimizing a combined loss function comprising categorical cross-entropy (classification task) and mean squared error for magnitude and for *ω* inferences (regression tasks), weighted 0.4, 0.3, and 0.3, respectively. We used the Adam optimizer (Kingma and Ba 2017) with an initial learning rate set by the hyperparameter search, reducing the learning rate by half when the loss on the early-stopping split failed to improve for 7 consecutive epochs (down to a minimum of 10^−6^), and stopped training when it failed to improve for 30 consecutive epochs, restoring the model weights from the best-performing epoch. We scored each hyperparameter combination in each fold using a composite metric combining shift-classification accuracy, the coefficient of determination (*R*^2^) for magnitude inference, and *R*^2^ for *ω* inference on the evaluation split, weighted 0.4, 0.3, and 0.3. We then averaged this score across the 10 folds (Table S1).

We selected the hyperparameter combination with the highest mean score (learning rate 10^−3^, dropout rate 0.2, dense-layer sizes 256 and 128) and retrained a single final model on the tuning set split into 12,150 matrices for training and 1,350 for validation (early stopping). Input and target standardization used the mean and standard deviation from this final training split, which we then applied to the held-out test set. We summarized model performance on the test set by classification accuracy, *R*^2^, and mean absolute error (MAE).

To test whether the trained CNN generalizes beyond the parameter combinations on which it was trained, we simulated six additional parameter combinations (gradual shift, magnitude = 40, and six values of *ω* used above). All other simulation parameters and the optimum trajectory were identical to those described in Methods: Quantitative trait simulation, and we ran 100 replicates per combination. Each replicate was summarized into a 20 × 40 genetic architecture matrix following the same procedure described in Methods: Deep learning and standardized with the mean and standard deviation of the final training set described above. Predictions were then made with the final model without further training.

### Use of artificial intelligence tools

ChatGPT (OpenAI) was used to assist with the algebraic derivation of the analytical results presented in the Supplementary Note. The authors supplied the standard expression for mean population fitness (Bürger and Lynch 1995, equation (3a)) together with the extinction threshold adopted here, and asked the tool to solve for the distance between the population mean phenotype and the new optimum at which genetic variance determines whether mean fitness falls below that threshold. The intermediate steps generated by the tool were used as a guide rather than as final results. Every step was independently rederived and verified by the authors, and the complete derivation was checked independently (see Acknowledgments). ChatGPT was also used to assist in writing the Python code for training the deep learning models used in this study. All code was reviewed, tested, and revised by the authors, and is available at https://github.com/YunaZhang73/simulate_scenarios. Claude (Anthropic) was used to suggest improvements to spelling, grammar, and wording in text written by the authors. All suggestions were reviewed and further revised by the authors. The final arguments, interpretation, and conclusions represent the authors’ own work and judgment. The authors take full responsibility for the content of this manuscript.

## Results

The evolution of a quantitative trait is a complex process that is influenced by the way the environment changes, the magnitude of environmental shift, and the genetic architecture of the trait. To understand if and how populations can adapt to the multitude of potential combinations of required trait adaptations to environmental change, we simulated polygenic traits and their response to a 400-generation period of environmental shift followed by a return to a constant environment. We compare across 150 parameter combinations defined by three types of phenotypic optimum shifts (sudden, gradual, and fluctuating shifts), a range of optimum shift magnitudes (amount of phenotypic change required for the population to reach the new phenotypic optimum), and different trait architectures, characterized by the size of the standard deviations of mutational effect sizes (*ω*).

First, we evaluated the adaptive trajectories of adaptation for a trait with intermediate effect sizes responding to a moderate environmental change (magnitude=5) during a sudden, gradual, or fluctuating shift. The type of environmental shift produced markedly divergent evolutionary trajectories of the phenotype. During a sudden environmental change, the mean phenotype increased most rapidly among the three shifts (Figure 1B) and the genetic variance increased alongside, before declining as the mean phenotype stabilized near the new phenotypic optimum (Figure 1C). While the mean fitness of the population dropped sharply to 0.29 immediately after the sudden shift, it rapidly recovered and exceeded 0.9 within 25 generations, leading to a successful adaptation (Figure 1C). However, when the magnitude of environmental shift increased, the population fitness was close to zero, implying that populations went extinct (Figure S4).

**Figure 1.**
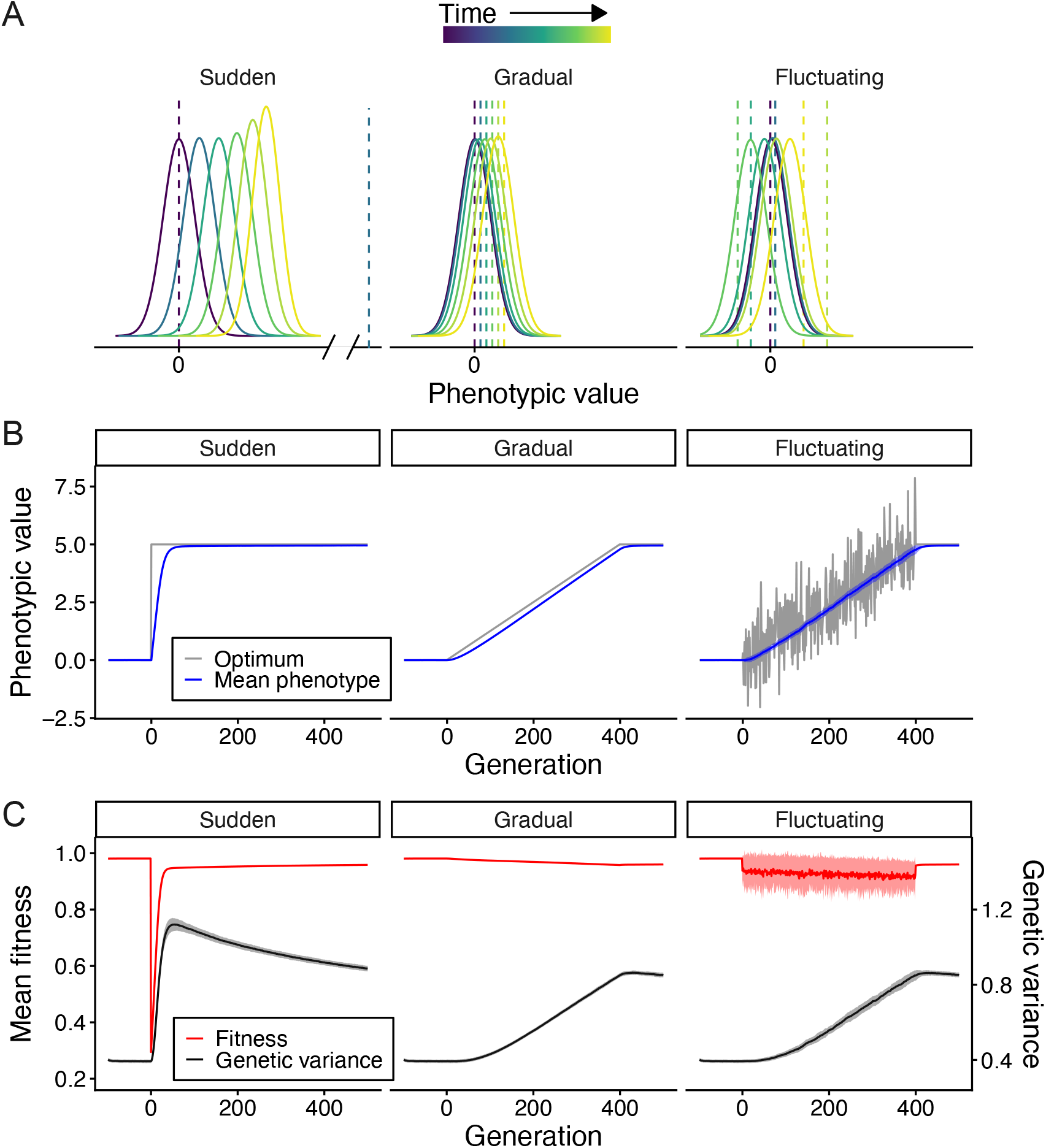
Polygenic adaptation to different environmental shifts. **(A)** Schematic illustration of phenotypic evolution over several generations during sudden, gradual, and fluctuating environmental changes. Solid curves represent phenotypic distributions, while dashed vertical lines denote the trait optimum. The color gradient (dark blue to yellow) indicates the progression of time, beginning with the initial state at zero (dark blue). During sudden change, the optimum shifts instantaneously to a new value. During gradual change, the optimum moves at a constant speed toward the new value. During fluctuating change, the optimum fluctuates stochastically while trending toward the new value at the speed of the gradual change. **(B)** Simulated phenotypic dynamics for a quantitative trait (*ω* = 0.3) across different types of environmental shift with a magnitude of 5 ( equal to a change of 0.02 Haldanes after adaptation). Gray lines represent the optimal phenotype (in the fluctuating panel, the line represents the trait optimum trajectory from a single simulation), and blue lines represent the mean of 100 independent mean phenotypes across simulation replicates. Shaded regions indicate the standard deviation (mean ± SD.). **(C)** Mean fitness (red) and genetic variance (black) for the same parameter combinations as in (B). Shaded regions indicate the standard deviation (mean ± SD.).

During gradual environmental change, the mean phenotype tracked the steadily moving optimum with a lag, which widened during the early response to selection and decreased as the mean phenotype caught up with the shifting optimum, before converging to the new optimum once the environment did not change anymore (Figure 1B). The fitness during gradual changes remained high and depended little on *ω* (Figure S4). Yet traits with larger *ω* initially decreased the mean phenotypic lag faster, resulting in even higher fitness compared to traits with smaller *ω* (Figure S5). Following an initial decline in fitness and partial recovery, the mean fitness continued to rise slowly for small-*ω* traits (*ω* = 0.05, 0.1), whereas it later decreased for larger-*ω* traits (Figure S6). The patterns reflect the complex interplay between the mean phenotypic lag and genetic variance (*V*_*G*_), as captured by the expression for mean fitness for normally distributed phenotypes (Equation (S1)), see equation (3a) of Bürger and Lynch (1995).

During fluctuating environmental shifts, the average responses of the mean phenotype, mean fitness, and the genetic variance were largely similar to those observed for the gradual shift, albeit with greater variation across generations and replicates (Figure 1B-C, Figures S4 and S5). Notably, despite the differences being small, random environmental fluctuations caused a lower mean fitness compared to those under the same parameter combinations during the gradual shifts. Moreover, the mean fitness of small-*ω* traits exhibited high variation across replicates, leading to occasional population extinctions (Figures 2 and S4). Hence, fluctuations had an impact on the adaptive success of a population.

**Figure 2.**
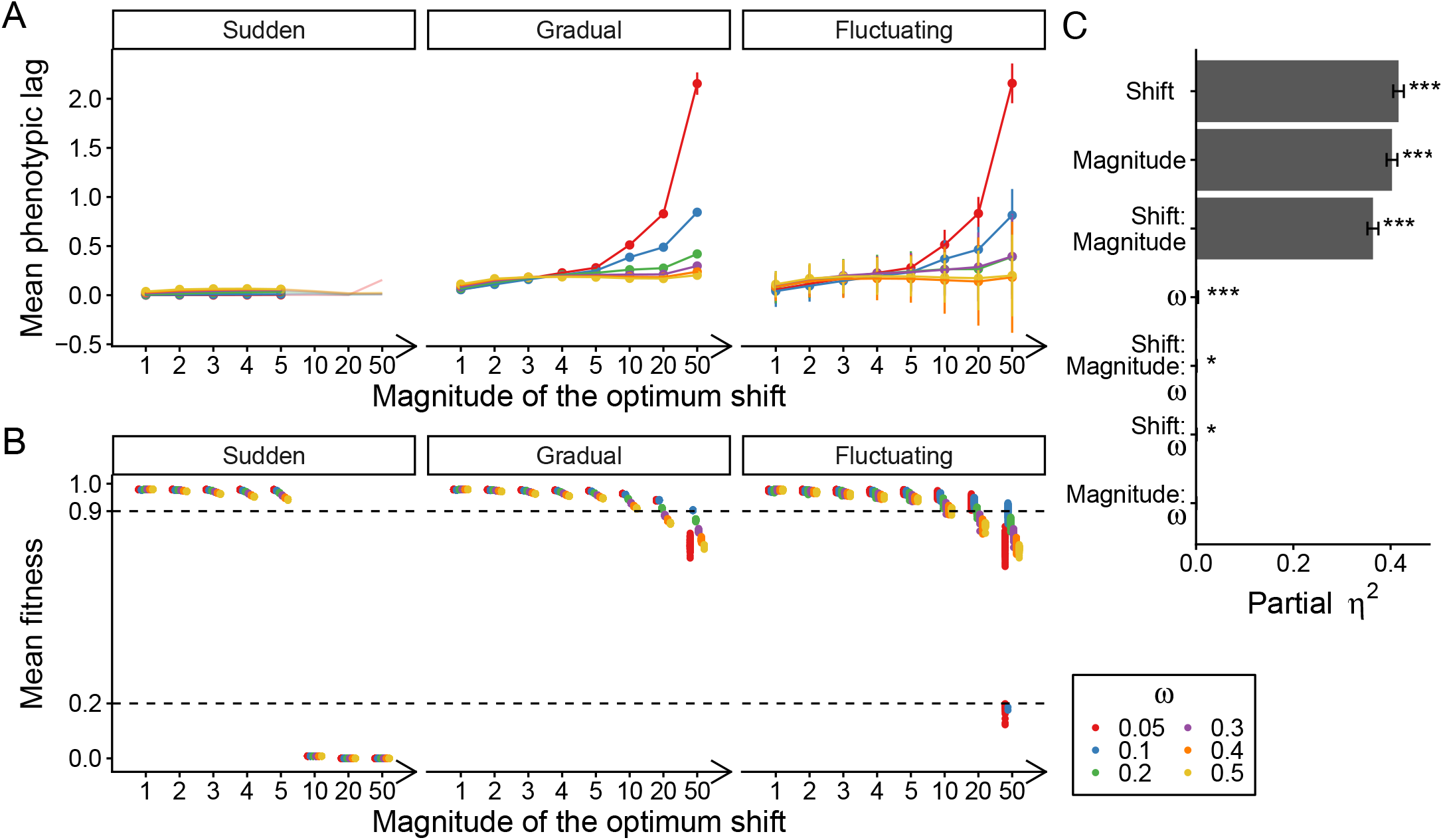
Evolutionary outcomes after 400 generations. **(A)** Mean phenotypic lag, describing the difference between the mean phenotype and the phenotypic optimum (averaged across 100 replicates). Colors describe the trait with a *ω* value at the three types of environmental shift (panel) for different magnitudes of shift (x-axis). The error bars represent the standard deviation of the mean lags across 100 replicates. Parameter combinations where more than 50% of populations went extinct are shown in lines with transparency. **(B)** Mean population fitness in each simulation (100 replicates per parameter set in total). Dashed horizontal lines indicate the fitness thresholds for adaptation (0.9) and extinction (0.2). **(C)** ANOVA results for mean fitness. Bars show the variance explained (partial *η*^2^) by the environmental shift, the magnitude of environmental shift, the effect size distribution of new mutations (*ω*), and their interactions. Error bars represent 95% confidence intervals, and the asterisks denote significant levels (* p<0.05; *** p<0.001).

### Evolutionary outcome and its determinants

The temporal dynamics of optimum shifts give an overview of the process of polygenic adaptation. To compare the long-term outcome of adaptation, we evaluated the population mean fitness 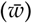 at generation 400 when the optimum across the three shifts had converged to the same value. Successful adaptation was defined as 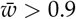 and extinction as 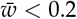. After sudden environmental shifts, populations either adapted successfully or immediately went extinct. While populations were able to adapt to less severe shifts (small magnitude), populations went extinct for magnitudes larger than or equal to 10, regardless of the trait architecture (Figure 2B and Figure S4). Gradual shifts, by contrast, did not lead to extinctions; however, successful adaptation depended on both the magnitude of the optimum shift and the *ω* value of the trait. After low-magnitude gradual shifts, all populations adapted successfully. After high-magnitude gradual shifts, maladaptation manifested in two distinct patterns: traits with small *ω* failed to track the moving optimum (*ω* = 0.05 at magnitude=50), whereas traits with large *ω* could approach the new optimum but suffered a fitness cost from elevated genetic variance, reducing their mean fitness at the end of the optimum shift (*ω* ≥ 0.3 at magnitude 20; and *ω* ≥ 0.2 at magnitude 50) (Figure 2B and Figure S7). Consequently, after gradual environmental shifts with a magnitude of 50, only traits with *ω* = 0.1 successfully adapted by generation 400, suggesting that effective adaptation requires an intermediate *ω* value that balances a reduction in phenotypic lag and genetic variance to maintain high fitness in the population. With additional fluctuations, maladaptation emerged at even lower magnitudes, starting at 10. More critically, severe fluctuations at magnitude 50 led to extinction in populations with *ω* = 0.05 and 0.1 (Figure 2B).

To quantify the effects of each input variable (shift, magnitude, and *ω*) and their interaction on evolutionary outcomes, we fit a linear model to explain fitness in generation 400 and estimate the effect size of each factor (partial *η*^2^). Within our simulation parameter space, the type of environmental shift had the largest effect on mean fitness, followed by the magnitude of the shift, with a substantial interaction between the two (Figure 2C). In contrast, the effect of *ω* was small, likely because its influence only manifested under high-magnitude shifts. Likewise, interactions involving *ω* accounted for little variance in mean fitness at generation 400 (Figure 2C). In summary, the type and magnitude of environmental shifts had the largest influence on the adaptive outcome for a population.

### Standing variation is the primary driver during polygenic adaptation

A population can adapt either through new mutations that arise after the trait optimum starts moving or through mutations that are already segregating in the population when the optimum starts moving. The relative contribution of each type of variation to successful adaptation might differ depending on the trait architecture and type and strength of environmental change. We quantified the phenotypic contribution of standing variation over the course of adaptation by assessing mutations and separating them by the time they arose.

Looking at the dynamics in an individual population (Figure 1B; magnitude=5, *ω* = 0.3) adapting to the three different types of optimum shifts provides intuition of the process. Standing variation primarily fueled the initial adaptive response across all environmental shifts, indicating its critical importance during the early stage of adaptation (Figure 3A). During sudden shifts, standing variation enabled rapid phenotypic change, minimizing the lag between the mean phenotype and the new optimum. During the subsequent fine-tuning phase, *de novo* mutations gradually increased in selective advantage, leading to a modest increase in their cumulative phenotypic contribution. During gradual and fluctuating shifts, the absolute contribution of standing variation increased over time, but its proportional share declined, remaining lower than during the sudden shift (Figure 3A and Figure S8).

**Figure 3.**
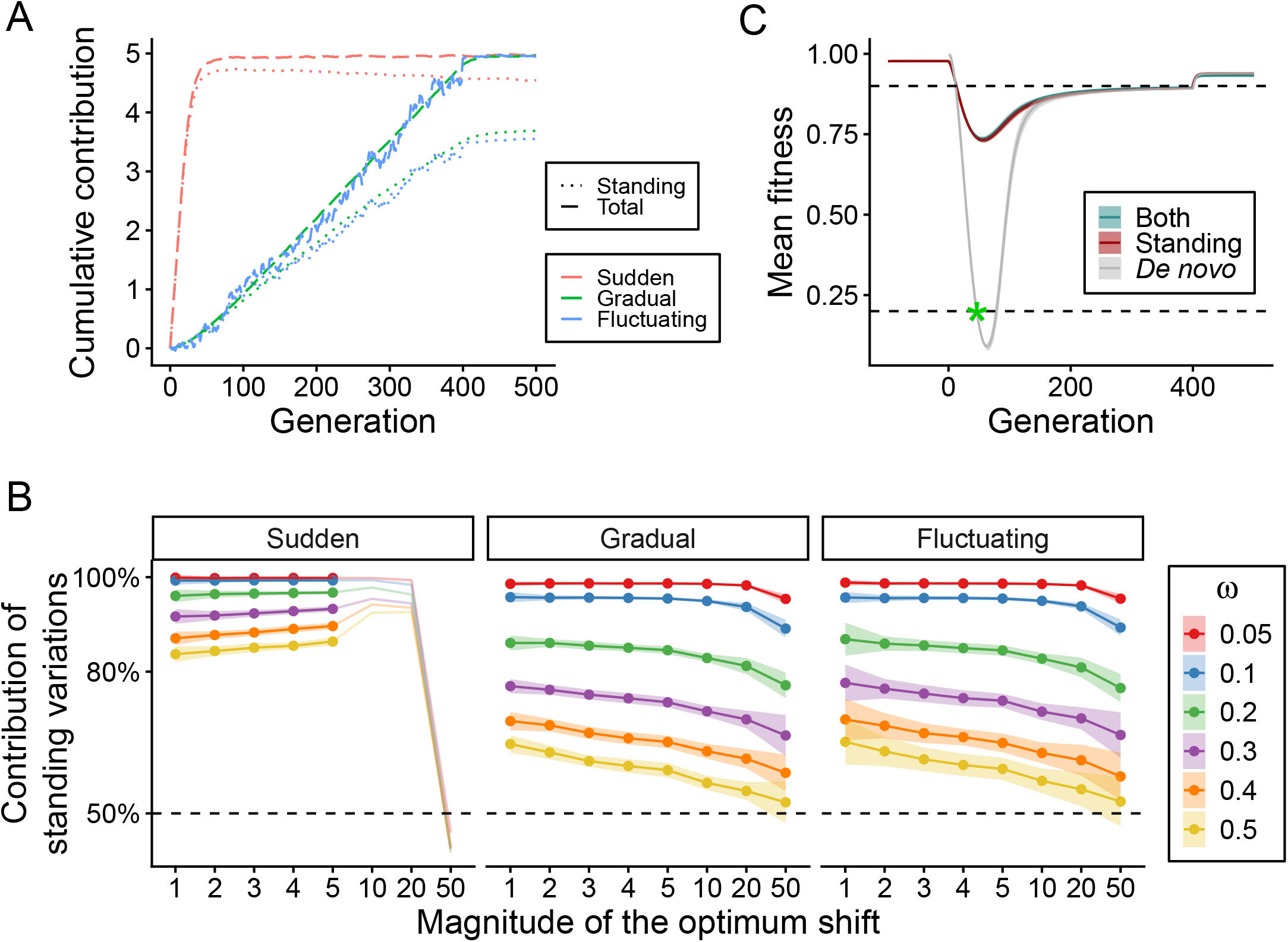
Standing variation drives adaptation. **(A)** Cumulative phenotypic changes contributed by standing variation over time. Colors denote shift types; dashed line represents the cumulative contribution of standing variation, solid line represents the contribution of all mutations (standing + *de novo* mutations). Results shown for one example trait (magnitude=5 and *ω* = 0.3). **(B)** The proportion of total phenotypic change after 400 generations that is attributed to mutations that pre-existed at generation 0 (standing variants). The dots represent the average proportion across 100 replicates, and the shaded areas indicate the standard deviation. The values of extinct populations are shown in lines with transparency. **(C)** The dynamics of mean fitness in simulations with only standing mutations (red), only *de novo* mutations (gray), or both (blue). Shaded regions represent the standard deviation across 100 replicates (mean ± SD.). Dashed horizontal lines indicate the fitness thresholds for adaptation (0.9) and extinction (0.2). The green asterisk denotes the occurrence of population extinction.

We then summarized the contribution of standing variation at the end of the environmental change (generation 400). We found that standing variation accounted for more than 50% of total phenotypic change across most parameter combinations. This dominance was most pronounced during sudden environmental shifts, where the contribution of standing variation exceeded 80%, underscoring the critical role of pre-existing genetic diversity in enabling rapid adaptation to abrupt environmental shifts (Figure 3B). We found a strong negative correlation between *ω* and the reliance on standing variation: under the same magnitude of environmental change, standing variation contributed more to phenotypic change in small-*ω* traits than in large-*ω* traits (Figure 3B).

Populations with larger genetic variance tolerate larger phenotypic lag than populations with lower genetic variance during sudden environmental shifts. Analytically, this holds true for our parameters as long as the extinction threshold of mean fitness is lower than 0.6 (see Supplementary Note for the derivation).

However, for gradual and fluctuating environmental change, the timing of the lowest mean fitness is less predictable. In addition to our simulations with standing and *de novo* mutations, we simulated populations evolving without *de novo* mutations, and populations evolving without standing variation for a parameter combination of *ω*=0.1 and a magnitude of 50 during a gradual shift. Populations lacking standing variation went extinct within tens of generations (Figure 3C). Conversely, populations without *de novo* mutations performed similarly to those with both sources of genetic variation (Figure 3C). These results demonstrate the indispensable role of standing variation in maintaining population persistence during rapid environmental change.

### Fixations have a limited contribution to polygenic adaptation

Given that standing variation plays a dominant and indispensable role in the adaptive process, it is important to understand how it facilitates adaptation. Positive-effect mutations could fix to advance the population, or a larger number of mutations change only mildly in frequency rather than fixing. In our simulations, the number of standing variants that were lost throughout the 400 generations was generally large across all combinations of parameters, while the number of fixed standing mutations was very low in most cases (Figures S9 and S10).Only when small-*ω* traits faced a high magnitude of change, the number of fixations significantly increased compared to traits in the static environment, suggesting a pronounced decrease in genetic diversity. However, despite this heterogeneity across traits, the combined contribution of both fixations and losses was less than 20% of the total phenotypic change for all surviving populations, indicating that most of the adaptive response was driven by segregating sites (Figure 4A).

**Figure 4.**
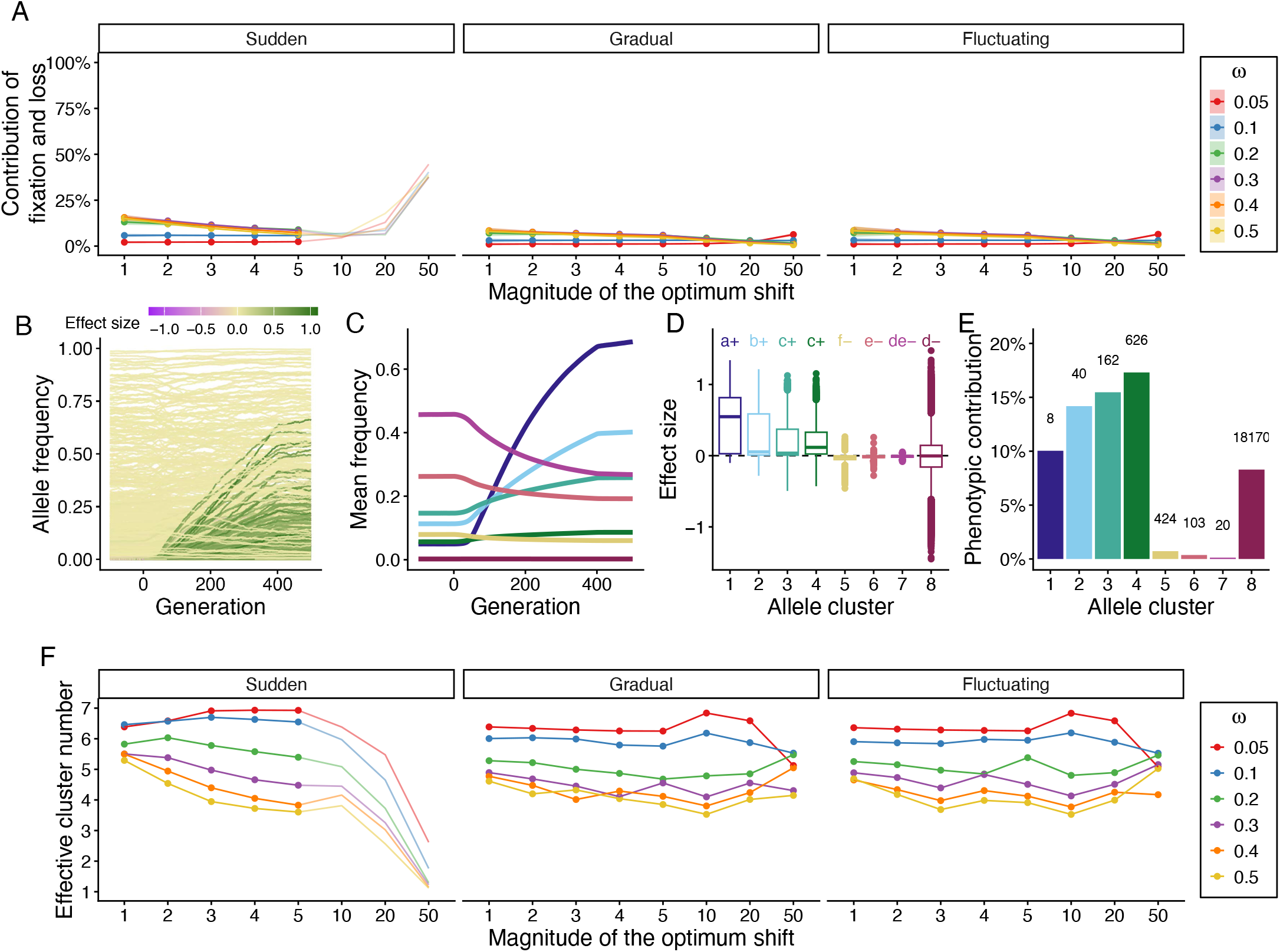
Allele frequency changes of standing variation. **(A)** Proportion of total phenotypic change after 400 generations of evolution caused by fixations and loss of standing mutations. The dots represent the average across 100 replicates; the shaded areas indicate the standard deviation. The proportions in extinct populations are shown in lines with transparency. Data shown in **B-E** are from the parameter combination of gradual shift, magnitude 50, and *ω* = 0.3. **(B)** Allele frequency trajectories of all standing variants from a single simulation. Each line is an individual allele. Colors represent the effect size of the mutation. **(C)** Mean allele frequency trajectories of the 8 clusters, identified by PCA and k-means clustering on allele frequency trajectories across 100 simulations. **(D)** Effect sizes of alleles in each cluster. Boxplots show median (center line), interquartile range (box), and whiskers extending to 1.5× interquartile range. Points represent outliers. Letters denote statistically significant differences among groups (ANOVA with post-hoc Tukey test, *P <* 0.05). The sign on the right side of the letter indicates the sign of the mean effect size. **(E)** Each bar represents the proportion of phenotypic change contributed by each cluster to the total change. The number above denotes the mean number of alleles in each cluster across the 100 simulations that were used to cluster. **(F)** Effective number of clusters out of the first seven examined clusters (cluster 8 was excluded) that contributed to adaptation across parameter combinations.

**Figure 5.**
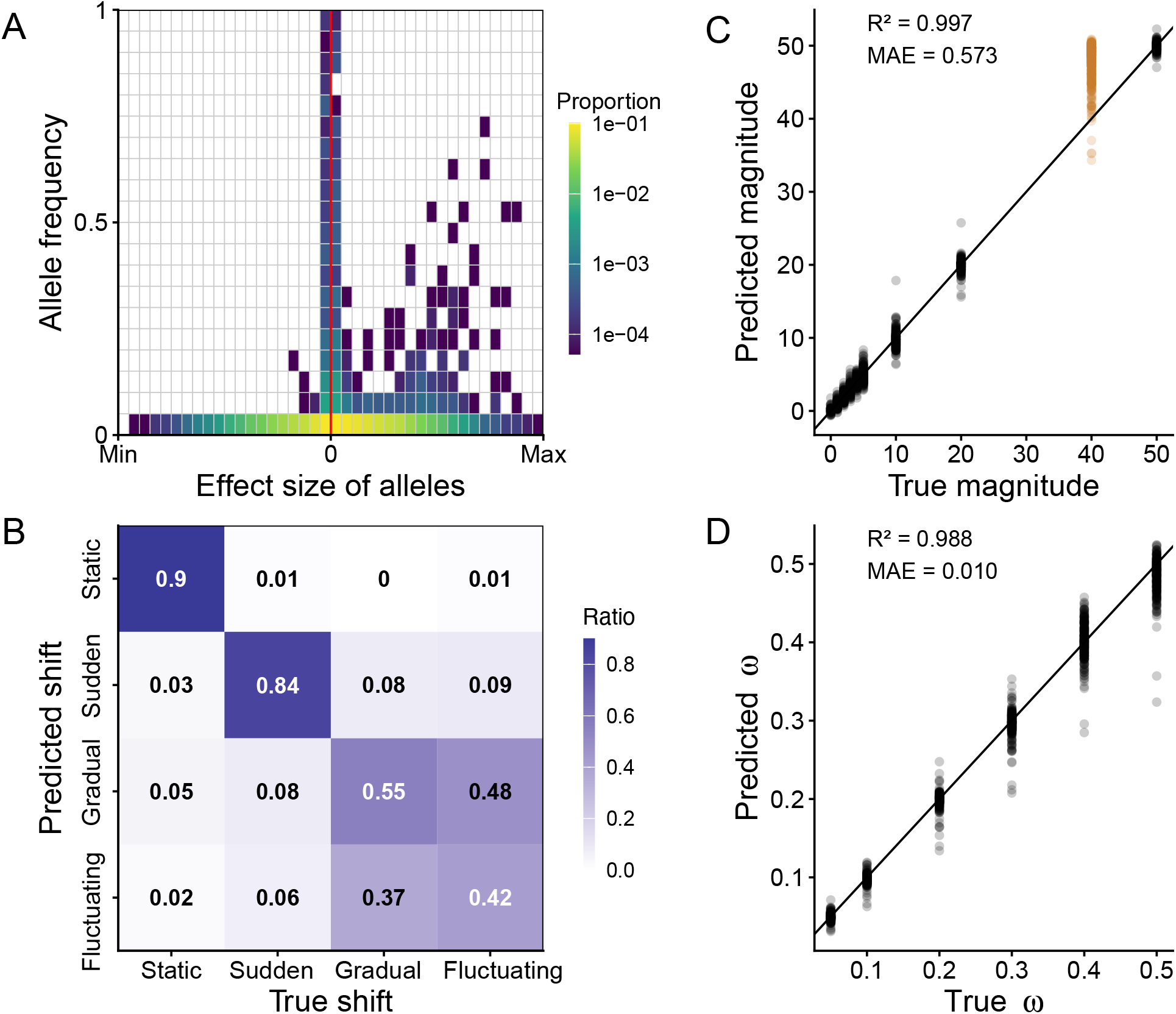
Parameter inference with Convolutional Neural Network (CNN). **(A)** An example of the genetic architecture matrix used as input for training the CNN. The matrix element represents the proportion (indicated by color) of alleles falling within specific ranges of effect sizes (columns) and allele frequencies (rows) in the population. The red vertical line separates negative and positive effect sizes. The data shown are from one simulation of a population with *ω* = 0.3 adapting to a gradual environmental change of magnitude 50. **(B)** Confusion matrix summarizing model performance in classifying environmental shift on the held-out test set. Each entry (denoted by color) represents the ratio of instances in which samples from a given true shift (x-axis) were predicted as a particular shift (y-axis). Diagonal entries correspond to correct classifications. **(C)** Results for inferring the magnitude of environmental changes on the held-out test set (black transparent dots) and on an unseen dataset of magnitude 40 (brown transparent dots). **(D)** Results for inferring the *ω* value on the held-out test set.

### Classification of allele frequency trajectories enables comparison of processes

To understand how adaptive genetic variants changed in frequency over time, we analyzed the frequency trajectories of all standing variants over the complete time of environmental change. The number of standing mutations per simulation replicate ranged from 16,000 to 32,000 per population (this number increased as *ω* decreased). To capture the major patterns from this high-dimensional dataset (Figure 4B), we performed a Principal Component Analysis (PCA) on the allele frequency-time matrix across all 100 replicates for each parameter combination. The first three PCs collectively explained more than 90% of total variance. We then applied a k-means clustering to assign the mutations to 8 clusters (see Figure S3 for the choice of *k*). Although the clusters did not occupy clearly separated regions in the PC space, indicating that allele frequency trajectories form a continuous spectrum rather than discrete classes (Figure S11), they nevertheless follow a trajectory pattern, summarizing the major trajectories of alleles.

The trajectories are illustrated for a single trait (gradual shift, magnitude=50, *ω* = 0.3), where visually reading the individual trajectories is not possible, but after applying our clustering, the first 7 clusters represent interpretable trajectories (Figure 4B-C). Alleles in cluster 8 remained at consistently low frequencies on average throughout the simulation (Figure 4C). We sorted the first seven clusters according to their mean change in frequency. Alleles in clusters 1-4 increased in frequency from generation 0 to 400 and had positive mean effect sizes (Figure 4C-D). Cluster 1 showed the mean frequency shifted from 0.05 at generation 0 to 0.67 at generation 400, and these alleles had significantly larger effect sizes than alleles in other clusters. More alleles with smaller effect sizes showed milder frequency increases (clusters 2-4), but contributed greater proportions to phenotypic change (Figure 4C-E). Cluster 4 was the largest contributor to phenotypic change, despite having small effects and small frequency changes. Yet, the number of mutations in this cluster was high (Figure 4E). Alleles in clusters 5-7 declined in frequency and had negative mean effect sizes (Figure 4C-D). Analogous to the positively selected alleles in cluster 1, cluster 7 showed the most pronounced frequency decline, but comprised fewer alleles than clusters 5 and 6 (Figure 4C and E). Although negative selection acted on a substantial number of alleles, its contribution to total phenotypic change was small during strong directional selection as observed in the example population. Overall, these results demonstrate that during a gradual environmental shift with a high magnitude, polygenic adaptation is achieved through the collective response of many genetic variants with diverse trajectories, rather than through a single dominant trajectory.

The summary of allele frequency trajectories enabled us to compare allele frequency changes across the large set of parameter combinations. We calculated the effective number of clusters required to produce the observed contribution. Cluster 8 was excluded because it was dominated by non-adaptive trajectories and some ambiguous assignments, which cannot represent a clear adaptive pattern. The effective number of clusters of the example shown above was 4, aligning with the result in Figure 4E. Across all surviving populations, the effective number of clusters was higher than 3, suggesting that adaptation to different environmental changes is polygenic and employs different trajectory classes (Figure 4F). During low to moderate shift magnitudes, the effective number of clusters was higher for smaller *ω*, indicating that clusters with declining allele frequency trajectories can also explain a substantial proportion of phenotypic adaptation. However, during very rapid (high magnitude) environmental shifts, the effective number of clusters converged to 4-5 across all *ω*, suggesting a dominance of positively selected alleles even for small-*ω* traits.

### Decoding population history via deep learning

A major challenge is to reconstruct the selection history of a population from contemporary genetic data. The fact that a large portion of phenotypic change is achieved by allele frequency shifts at mutations that stay polymorphic in the population might allow the reconstruction of the selective history. We used deep learning to predict input parameters from the genetic architecture after environmental change. We summarized the genetic architecture as a 2D histogram of frequencies and effect sizes of segregating alleles at generation 400 (Stetter *et al*. 2018). We used a 20 × 40 matrix (Figure 5A) as input for a two-dimensional convolutional neural network (2D CNN). The CNN jointly inferred the environmental shift (classification task), the magnitude of this shift (regression task), and the standard deviation of the mutational effect sizes (*ω*; regression task). We selected model hyperparameters by grid search with 10-fold cross-validation on the tuning set (Table S1). Across the 32 hyperparameter combinations, the cross-validation performance of the two regression tasks was consistently high, whereas the classification accuracy for the environmental shift did not exceed 0.7, suggesting an inherent genetic similarity between some environmental shifts. We then retrained the final model with the selected hyperparameters on the tuning set, where the model achieved a robust balance between fitting the training data and maintaining generalization performance on the validation set (Figure S12).

Finally, we evaluated this model on the held-out test set. On this test set, the CNN demonstrated high power to identify static environments (90% accuracy) and sudden environmental shifts (84% accuracy) (Figure 5B). It also performed well in distinguishing gradual and fluctuating shifts from the other types, but was not able to confidently distinguish the gradual shifts from fluctuating shifts (Figure 5B). This reciprocal misclassification confirms that gradual and fluctuating environmental shifts produce similar genomic signatures. The CNN showed high predictive accuracy for both regression tasks. The predictive *R*^2^ for the magnitude of environmental shifts reached 0.997 with a MAE of 0.573, and the predictive *R*^2^ for *ω* achieved 0.988 with a MAE of 0.010 (Figure 5C-D).

To test the model performance on unseen parameters, we simulated an additional parameter set (gradual shift, magnitude = 40, and six values of *ω* used above) and applied the fitted model to the genetic architecture matrices extracted from these simulation data. This magnitude was not among the eight levels represented in the training data. Inferences of shift type and *ω* were unaffected (Figures S13 and S14), indicating that these inferences do not depend on the magnitude having been seen during training. Magnitude inference returned a mean of 48.3 with a standard deviation of 2.2 (Figure 5C). Despite the overestimation, the inferred results remain within the same order of magnitude as the true values.

Collectively, these results demonstrate that a 2D CNN can reveal the environmental history and trait architecture of a population from a single contemporary genetic snapshot.

## Discussion

The response to environmental change depends jointly on the environmental factors themselves, manifested by the type and magnitude of change, and on the traits that are under selection. To understand the adaptive process of quantitative traits driven by directional selection imposed by rapid environmental changes similar to those observed over recent centuries, we simulated a range of traits with different genetic architectures and subjected them to three types of optimum shifts of varying magnitudes. The optimum shifts we studied here ranged from mild changes 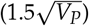 to strong changes 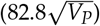, corresponding to 0.004 - 0.207 Haldanes for gradual and fluctuating shifts, while sudden shifts imposed the same change instantaneously. Similar rates of change have been documented for the advancement of parturition date in red deer adapting to sustained climate warming (≈ 0.02 Haldanes; Bonnet *et al*. 2019). However, the rates of phenotypic change documented in contemporary wild populations are predominantly low, with a long tail of fast values (Hendry and Kinnison 1999; Kinnison and Hendry 2001; Sanderson *et al*. 2022), and the sustainable rates are usually lower than 0.1 Haldanes (Kopp and Matuszewski 2014). The strong changes, by contrast, tend to be fleeting and arise from extreme environmental conditions or anthropogenic disturbance (Hendry *et al*. 2008; Sanderson *et al*. 2022), such as the change in beak size of Darwin’s finches during a single drought (Grant and Grant 2006), or the rapid evolution of shorter beaks in North American soapberry bugs tracking the smaller fruit of an invasive host plant (Carroll *et al*. 1997). Yet, due to the accelerating shifts through climate change, many populations will be faced with rates that require even higher trait adaptation rates. Shifts will manifest differently in different areas of the globe, and while some will be gradual with a small magnitude, others will appear as sudden shifts in trait optima in numerous species.

We compared the adaptive success of different trait architectures across three different types of optimum shift with a range of magnitudes, and found that both the type of shift and the magnitude had a substantial impact on the adaptive outcome. All populations adapted successfully to a low-magnitude change, and maladaptation emerged only as the magnitude increased, and extinction mostly occurred during sudden shifts of high magnitudes (Figure 2), indicating that mitigating environmental change would broaden the set of populations able to keep pace. During rapid gradual and fluctuating shifts (magnitude 50), adaptive success was governed by a trade-off set by *ω*. Larger *ω* generated more genetic variance during adaptation, which allowed the population mean phenotype to approach the moving optimum but also broadened the phenotypic distribution around it and decreased mean fitness (Figures 2 and S6). Populations with small *ω*, in contrast, failed to track the optimum, and the elevated phenotypic lag can ultimately lead to extinction (Figure 2; Bürger and Lynch 1995; Pahujani *et al*. 2026). Mean fitness was consequently highest at intermediate *ω* (Figure 2). Despite carrying a fitness cost in dispersion, the high genetic variance might not represent a disadvantage because it can provide a rich reservoir of variation that could facilitate adaptation to subsequent directional changes (Barrett and Schluter 2008) or avoid extinction (Figure 2). Specifically, large-effect adaptive alleles that remain in the population may become beneficial for future directional changes, while such alleles are rapidly lost or fixed during sudden shifts (Figure S15; Stetter *et al*. 2018). During fluctuating shifts, these large-effect alleles did not rise monotonically, but showed cross-generational frequency fluctuations superimposed on their directional trend (Figure S15). Similar fluctuations at many loci have been documented in natural populations during recurrent selection (Bergland *et al*. 2014; Kelly 2022) and in simulations of periodically changing optimum (Tuyishimire *et al*. 2026). The interplay of the trait architecture (*ω*), the type and magnitude of change thus governs whether populations persist in changing environments. Hence, decoding the genetic architecture of traits that matter for adaptation will be essential for predicting how species respond to rapid change.

Although the population genetic narrative has long suggested a prominent role for *de novo* mutations that arise after environmental change and then go to fixation to increase the fitness of a population, quantitative genetic results on trait architectures of adapted populations were not consistent with this view (Pritchard *et al*. 2010; Boyle *et al*. 2017; Sella and Barton 2019; Barghi *et al*. 2020). Yet, large-effect mutations have been found to leave signatures of selective sweeps (Bersaglieri *et al*. 2004; Fulgione *et al*. 2022). The increasing amount of genomic data from divergently adapted populations shows that even loci with large trait effects were often already segregating in the population before adaptation (Fairbanks and Ross-Ibarra 2025; Battlay *et al*. 2025; Wu *et al*. 2026). While the observations seem contradictory at first sight, large-effect mutations remain at very low frequency under stabilizing selection and are therefore less likely to be detected in sample sizes of only a few hundred (Sella and Barton 2019). Nonetheless, standing variation does not require waiting time to arise, and is therefore immediately accessible once selection pressure is imposed—an advantage that is especially critical during environmental shifts, where phenotypic change must be rapid (Figure 3B; Barrett and Schluter 2008). Furthermore, the higher starting frequency of standing variation not only increases its probability of being selected but also permits adaptation through two mechanisms: increasing the frequency of beneficial alleles and decreasing the frequency of deleterious alleles (Figure 4C-E). In our simulations, *de novo* mutations often contributed only a small portion of adaptive trait change, but individual mutations with large effects still rose to high frequency (Figures 3 and 4). Some of these loci would likely be identified as selective sweeps in empirical work. The moving optimum studied here strongly favored standing variation with smaller frequency shifts, but other selection modes, including truncation selection, leave greater scope for *de novo* mutations, once standing variation is depleted (Stetter *et al*. 2018). Our simulations show that standing variation remains available even during rapid change, particularly when change is gradual or fluctuating (Figure 3B). Allele frequency trajectories varied widely, from strong to weak frequency shifts, yet their contributions to phenotypic change were relatively evenly distributed, demonstrating that adaptation of quantitative traits is the collective result of many loci acting through varying degrees of frequency shifts and effect sizes (Figure 4F; Höllinger *et al*. 2019). The dispersed contributions of many loci became concentrated in fewer loci as the magnitude of change increased or when the mutational effect size distribution widened (large *ω*; Figure 4F).

Following allele trajectories in empirical data can be enabled through our clustering approach, which effectively grouped alleles with similar trajectories and similar effect sizes. As effect sizes are rarely available, but allele frequency trajectories become feasible for many species (Wu *et al*. 2026; Czorlich *et al*. 2018; Kelly 2022; Chen *et al*. 2019), our results suggest that we can infer effect size categories from trajectory clusters. Once effect size information is available, for instance from genome-wide association studies and genome-wide prediction models (Visscher *et al*. 2017; Josephs *et al*. 2017), the genetic architecture matrices can be used to infer the population’s selection history. The divergent adaptive genetic responses across environmental shifts imply that the resulting genetic architectures differ systematically at any given time point. Previous works have qualitatively described how the rate of gradual environmental shifts shapes genetic architecture (Pahujani *et al*. 2026), and a random forest model inferred *ω* from genetic architectures following a sudden environmental shift (Stetter *et al*. 2018). Here, we expand this framework using state-of-the-art machine learning models, by treating genetic architecture as two-dimensional, image-like data and training a CNN to jointly infer environmental shift, magnitude of the shift, and *ω*, from a single time-point snapshot after the adaptive process. Our CNN achieved high predictive accuracy, demonstrating that the abstracted genetic architecture retains sufficient signal to recover key aspects of selection history: the *ω* of the focus trait, the type of optimal phenotype shift induced by environmental shift, and the magnitude of that shift. Applying these models to environmental parameters outside the range of the training datasets or even empirical data has the potential to reconstruct selection histories from individual time points.

Collectively, our study advances the understanding of polygenic adaptation across a wide range of environmental contexts, provides a potential approach for inferring the population history from existing genetic data, and also provides valuable insights for conservation biology.

## Data availability

Code and data used for this study can be found at https://github.com/YunaZhang73/simulate_scenarios.

## Acknowledgments

We thank Sakshi Pahujani for carefully checking the mathematical derivations in this manuscript. We thank the members of TRR 341 *Plant Ecological Genetics* for insightful comments and discussions.

## Supplementary material

### Supplementary Note. Standing variation is beneficial under sudden environmental shifts

Assuming a sudden optimum shift is imposed on an equilibrium population under stabilizing selection. Selection acts on a phenotype through the Gaussian fitness function Equation (1). Thus, for an equilibrium population with a Gaussian phenotype, the population mean fitness is (Bürger and Lynch 1995, equation (3a)):

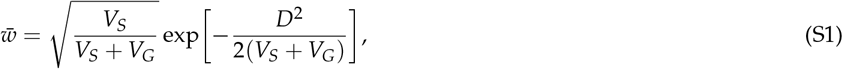

where 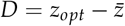 represents the mean phenotypic lag between the new optimum *z*_*opt*_ and the mean phenotype 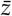. For a population with no standing variation (i.e., *V*_*G*_ = 0),

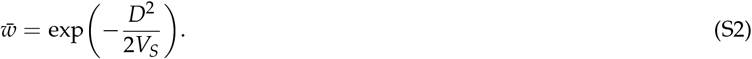

Let *D*_*vg*_ and *D*_0_ denote the maximum mean phenotypic lag that populations can tolerate before extinction, for populations with *V*_*G*_ *>* 0 and *V*_*G*_ = 0, respectively. Therefore, for a extinction threshold of fitness, *w*_*th*_, below which populations go extinct, we have

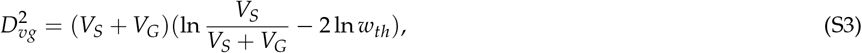

and

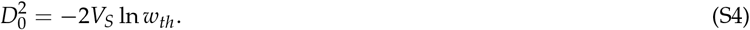

Hence, determining whether standing variation is beneficial under sudden environmental shifts reduces to asking whether *D*_*vg*_ *> D*_0_ for a given extinction threshold.

Since both *D*_*vg*_ and *D*_0_ are nonnegative, 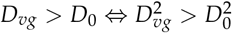. Substituting the expression for 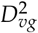 and 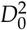 yields

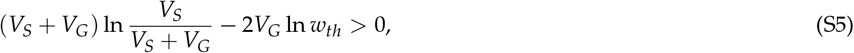

which can be rearranged as

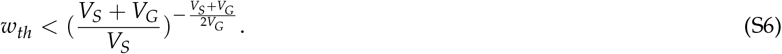

Let 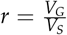, Equation (S6) can be rewritten as:

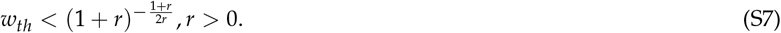

Let

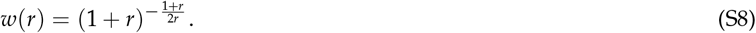

Taking logarithms,

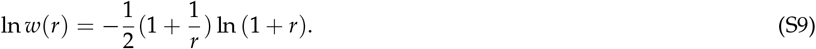

Take the limit, lim_*r* → 0_ *w*(*r*) = *e*^−1/2^ ≈ 0.607; lim_*r* → ∞_ *w*(*r*) = *e*^−∞^ = 0.

Taking derivatives,

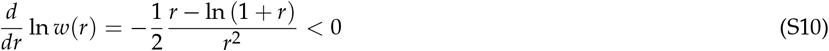

because ln (1 + *r*) *< r* for all *r >* 0. Hence, *w*(*r*) decreases monotonically with *r* and *w*(*r*) ∈ (0, 0.607). For small *r* (i.e., small *V*_*G*_), standing variation increases the maximum phenotypic lag that populations can tolerate as long as *w*_*th*_ is roughly less than 0.607. The upper bound of the extinction threshold fitness decreases as *V*_*G*_ increases.

In our simulations, the maximum standing genetic variance at generation 0 is *V*_*G*_ = 0.47, and *V*_*S*_ = 10. Therefore, standing variation is predicted to promote population persistence under sudden environmental shifts whenever *w*_*th*_ *<* 0.6. The extinction threshold fitness used in our analysis, *w*_*th*_ = 0.2, is well below this bound.

## Supplementary Table

**Table S1.** Results of hyperparameter grid search and 10-fold cross-validation. Mean composite score and its components (classification accuracy, *ω* regression *R*^2^, magnitude regression *R*^2^) averaged across 10 folds for each of the 32 hyperparameter combinations. The table is sorted in descending order by the mean score.

| Hyperparameters | Accuracy | R2_omega | R2_magnitude | Score |
| --- | --- | --- | --- | --- |
| {"dense1": 256, "dense2": 128, "dropout": 0.2, "lr": 0.001} | 0.631 | 0.989 | 0.996 | 0.848 |
| {"dense1": 256, "dense2": 128, "dropout": 0.5, "lr": 0.001} | 0.629 | 0.990 | 0.996 | 0.847 |
| {"dense1": 256, "dense2": 128, "dropout": 0.3, "lr": 0.001} | 0.629 | 0.988 | 0.995 | 0.847 |
| {"dense1": 256, "dense2": 128, "dropout": 0.2, "lr": 0.01} | 0.627 | 0.989 | 0.996 | 0.846 |
| {"dense1": 256, "dense2": 128, "dropout": 0.3, "lr": 0.01} | 0.626 | 0.990 | 0.996 | 0.846 |
| {"dense1": 128, "dense2": 64, "dropout": 0.2, "lr": 0.01} | 0.626 | 0.989 | 0.996 | 0.846 |
| {"dense1": 256, "dense2": 128, "dropout": 0.3, "lr": 0.0005} | 0.628 | 0.986 | 0.995 | 0.846 |
| {"dense1": 128, "dense2": 64, "dropout": 0.3, "lr": 0.001} | 0.626 | 0.988 | 0.995 | 0.845 |
| {"dense1": 256, "dense2": 128, "dropout": 0.4, "lr": 0.001} | 0.624 | 0.989 | 0.996 | 0.845 |
| {"dense1": 256, "dense2": 128, "dropout": 0.4, "lr": 0.01} | 0.623 | 0.989 | 0.996 | 0.845 |
| {"dense1": 256, "dense2": 128, "dropout": 0.5, "lr": 0.0005} | 0.626 | 0.987 | 0.993 | 0.845 |
| {"dense1": 256, "dense2": 128, "dropout": 0.2, "lr": 0.0005} | 0.624 | 0.987 | 0.994 | 0.844 |
| {"dense1": 128, "dense2": 64, "dropout": 0.3, "lr": 0.01} | 0.621 | 0.989 | 0.996 | 0.844 |
| {"dense1": 256, "dense2": 128, "dropout": 0.4, "lr": 0.0005} | 0.622 | 0.988 | 0.995 | 0.844 |
| {"dense1": 128, "dense2": 64, "dropout": 0.4, "lr": 0.001} | 0.622 | 0.988 | 0.993 | 0.843 |
| {"dense1": 256, "dense2": 128, "dropout": 0.5, "lr": 0.01} | 0.621 | 0.988 | 0.996 | 0.843 |
| {"dense1": 128, "dense2": 64, "dropout": 0.3, "lr": 0.0005} | 0.623 | 0.987 | 0.993 | 0.843 |
| {"dense1": 128, "dense2": 64, "dropout": 0.2, "lr": 0.001} | 0.621 | 0.987 | 0.995 | 0.843 |
| {"dense1": 128, "dense2": 64, "dropout": 0.2, "lr": 0.0005} | 0.620 | 0.986 | 0.995 | 0.842 |
| {"dense1": 256, "dense2": 128, "dropout": 0.2, "lr": 0.0001} | 0.621 | 0.985 | 0.993 | 0.842 |
| {"dense1": 128, "dense2": 64, "dropout": 0.4, "lr": 0.01} | 0.617 | 0.989 | 0.995 | 0.842 |
| {"dense1": 256, "dense2": 128, "dropout": 0.4, "lr": 0.0001} | 0.621 | 0.985 | 0.992 | 0.841 |
| {"dense1": 256, "dense2": 128, "dropout": 0.3, "lr": 0.0001} | 0.619 | 0.985 | 0.993 | 0.841 |
| {"dense1": 128, "dense2": 64, "dropout": 0.5, "lr": 0.01} | 0.616 | 0.985 | 0.994 | 0.840 |
| {"dense1": 128, "dense2": 64, "dropout": 0.4, "lr": 0.0005} | 0.617 | 0.987 | 0.990 | 0.840 |
| {"dense1": 128, "dense2": 64, "dropout": 0.2, "lr": 0.0001} | 0.615 | 0.985 | 0.991 | 0.839 |
| {"dense1": 128, "dense2": 64, "dropout": 0.3, "lr": 0.0001} | 0.615 | 0.985 | 0.992 | 0.839 |
| {"dense1": 256, "dense2": 128, "dropout": 0.5, "lr": 0.0001} | 0.614 | 0.984 | 0.991 | 0.838 |
| {"dense1": 128, "dense2": 64, "dropout": 0.5, "lr": 0.001} | 0.612 | 0.988 | 0.984 | 0.837 |
| {"dense1": 128, "dense2": 64, "dropout": 0.4, "lr": 0.0001} | 0.611 | 0.983 | 0.990 | 0.837 |
| {"dense1": 128, "dense2": 64, "dropout": 0.5, "lr": 0.0005} | 0.611 | 0.985 | 0.983 | 0.835 |
| {"dense1": 128, "dense2": 64, "dropout": 0.5, "lr": 0.0001} | 0.603 | 0.982 | 0.986 | 0.831 |

## Supplementary Figures

**Figure S1.**
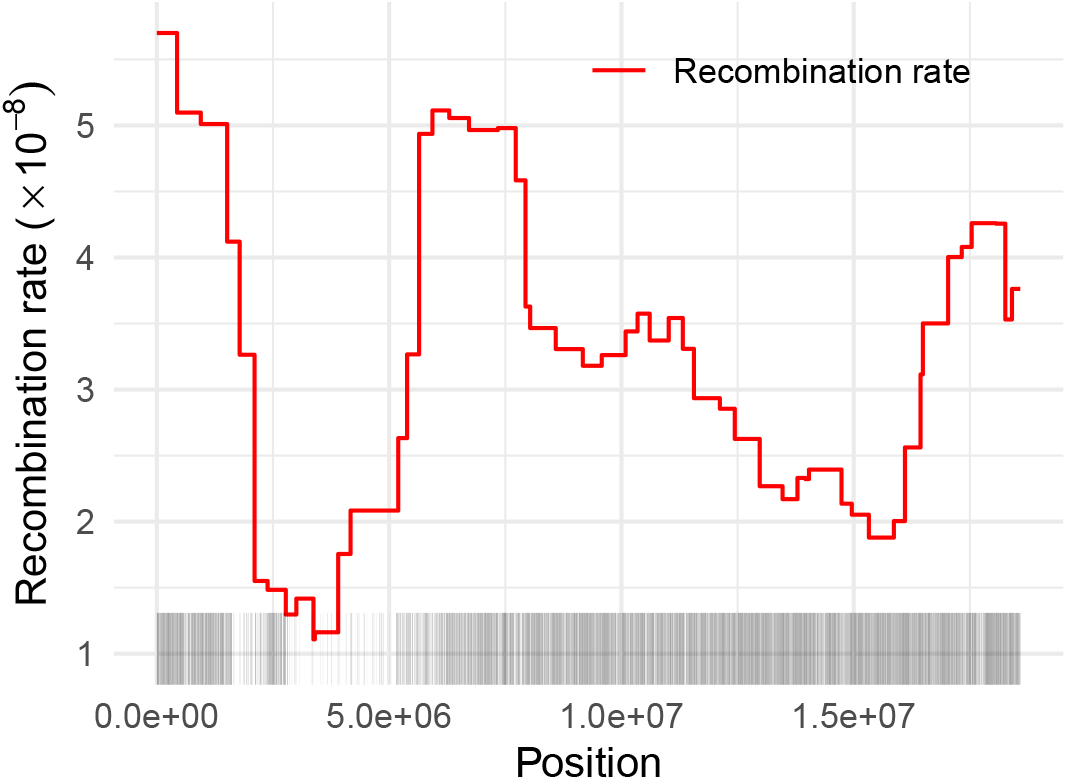
Genic region and recombination map. The lower gray lines denote the genic regions on chromosome 4 in *A. thaliana*. The upper red line indicates the recombination rate along the chromosome.

**Figure S2.**
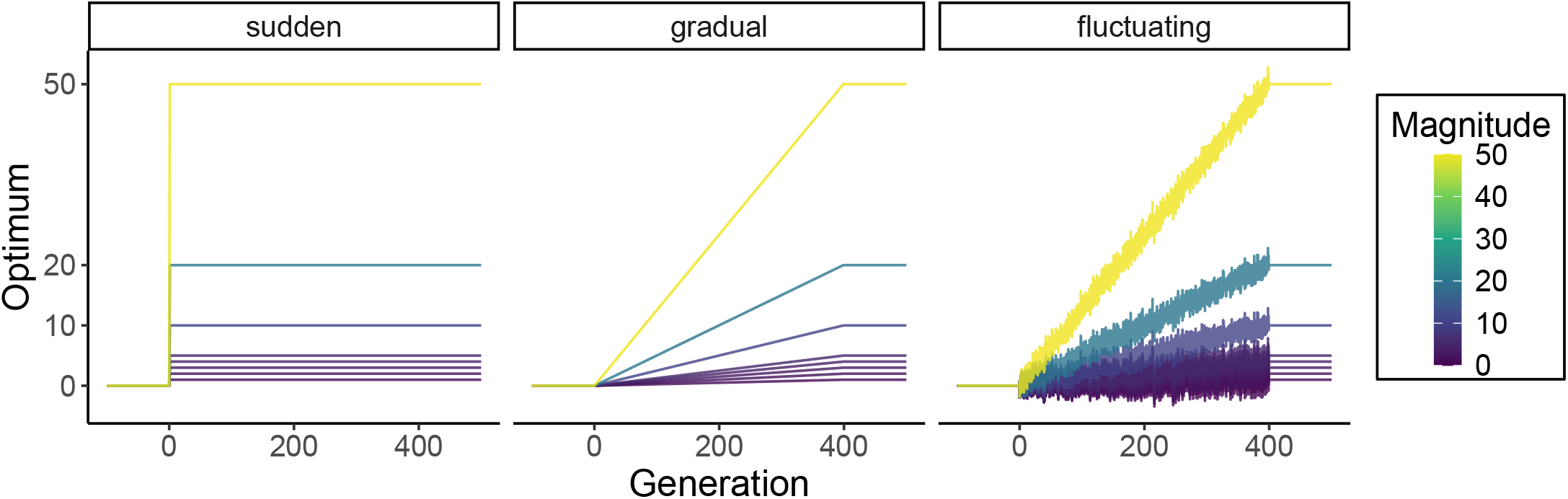
Trait optimum set for simulations. The trait optimum over the 600 generations (generation -99 to 500) following the 10*N* burn-in period. Each line represents a parameter combination of the mode and the magnitude of the trait optimum shift for the six different *ω* values used for simulations.

**Figure S3.**
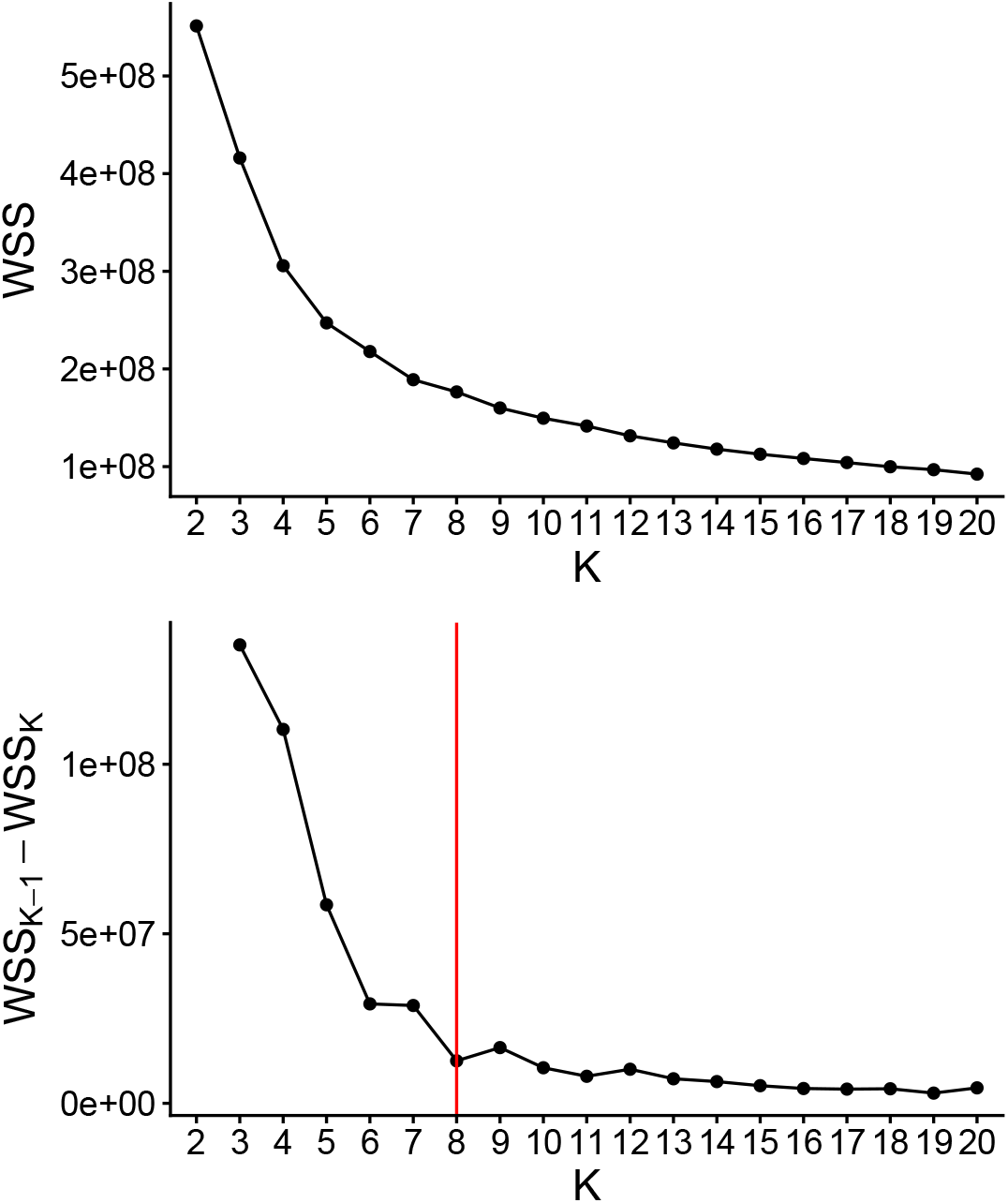
Selection of k value. The upper plot shows the total sum of squares within clusters (WSS) from the clustering results with different k. The bottom plot shows the reduction in WSS as k increases by one. Analysis was done on data from 100 replicates of the parameter combination: gradual shift, magnitude = 50, and *ω* = 0.3.

**Figure S4.**
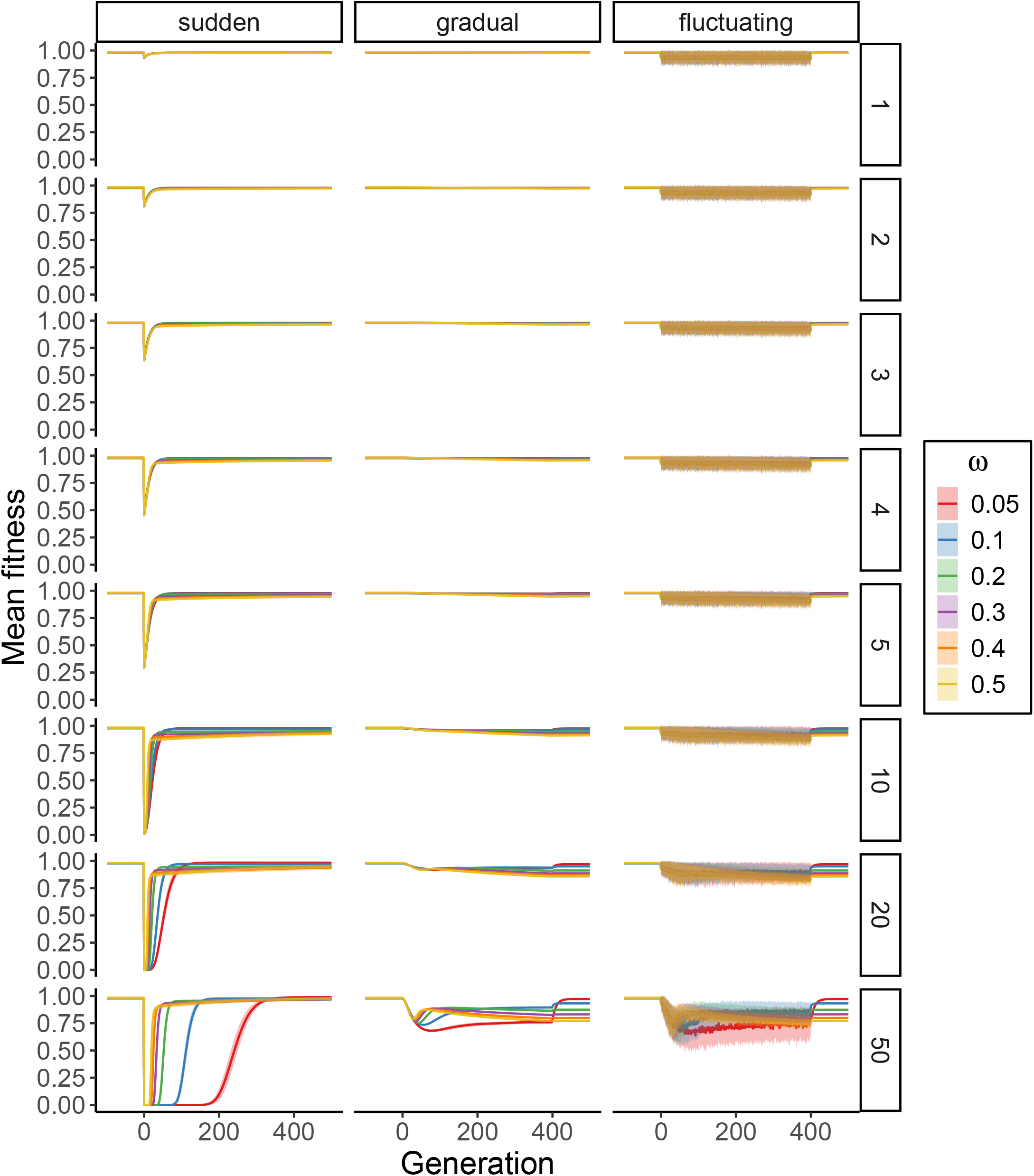
Mean fitness over time for different traits. Each panel represents the shift (column) and the magnitude (row) of environmental change. Each line shows the mean fitness across the individuals of 100 replicate populations, with the shaded region indicating the standard deviation across replicates’ means (mean ± SD.). Colors denote the value of *ω*.

**Figure S5.**
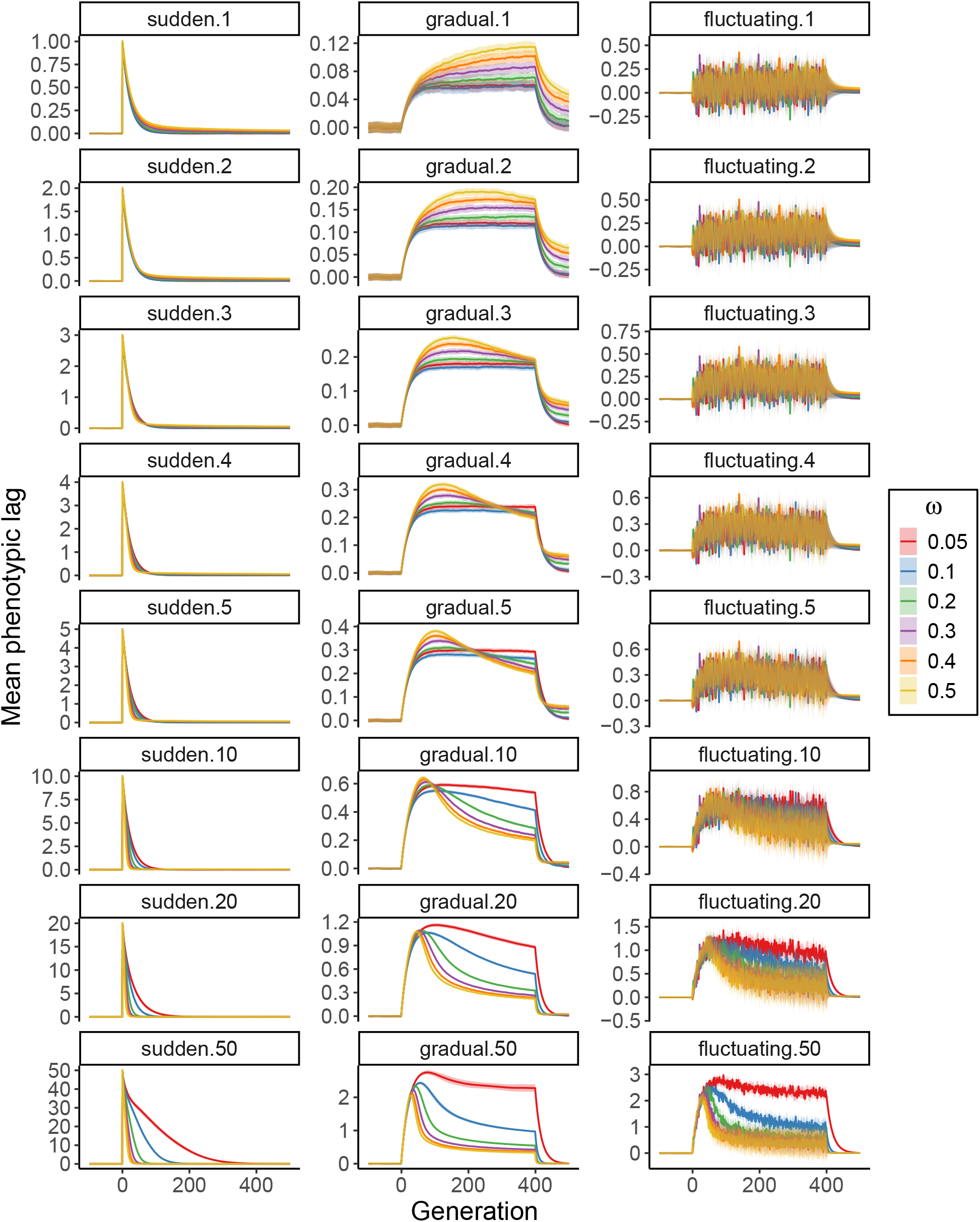
Mean phenotypic lag over time for different traits. Each panel represents the same environmental condition with a certain optimum shift and its magnitude, as indicated in the title of the panel ([shift].[magnitude]). Each line shows the mean phenotypic lag across the individuals of 100 replicate populations, with the shaded region indicating the standard deviation across the replicates’ means (mean ± SD.). Colors denote the value of *ω*.

**Figure S6.**
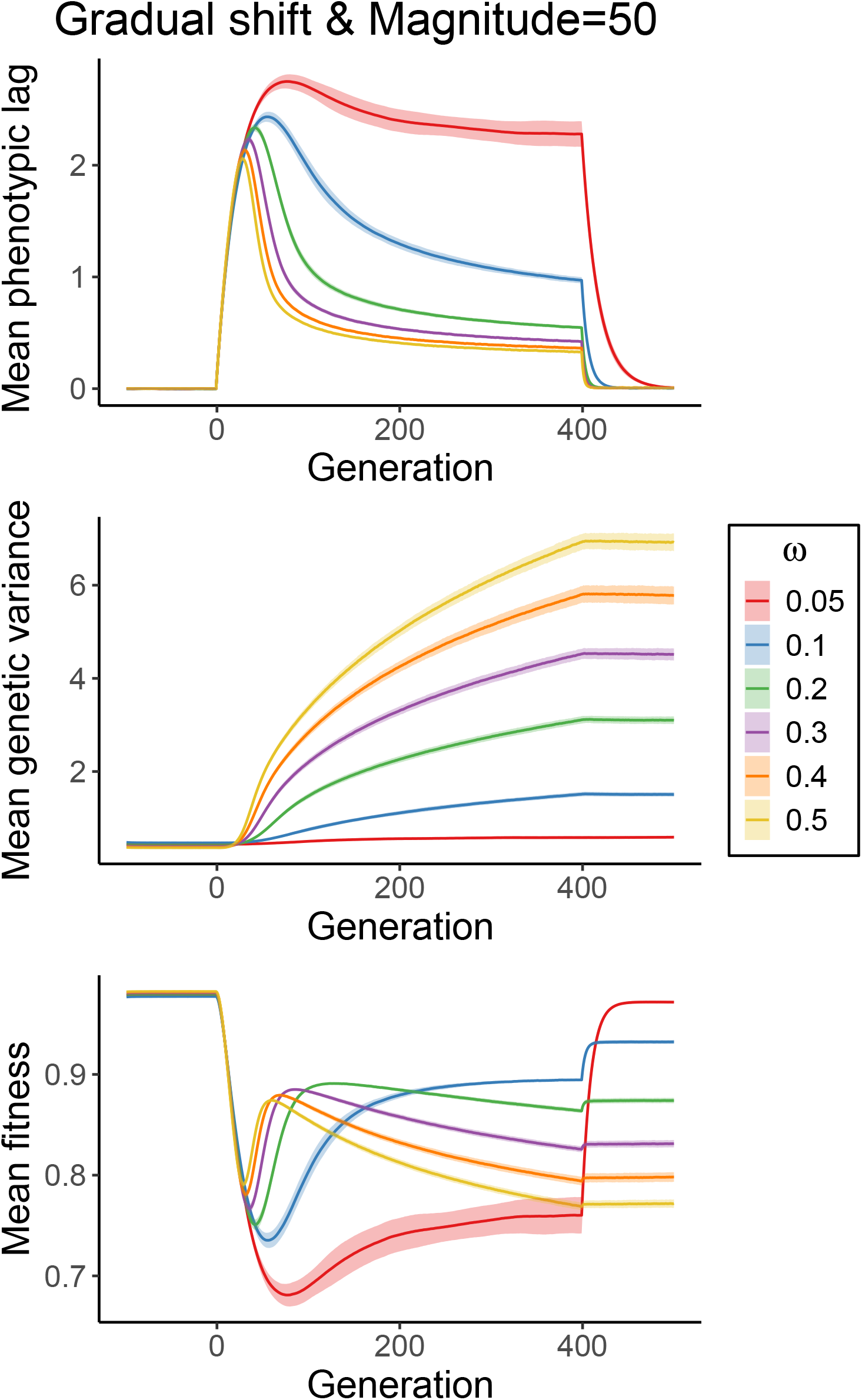
Population outcomes under gradual environmental change with a magnitude of 50. Mean phenotypic lag, genetic variance, and mean fitness of the populations over time. Each line represents the mean across replicates, with the surrounding shaded region indicating the standard deviation (mean ± SD.). Colors denote the value of *ω*.

**Figure S7.**
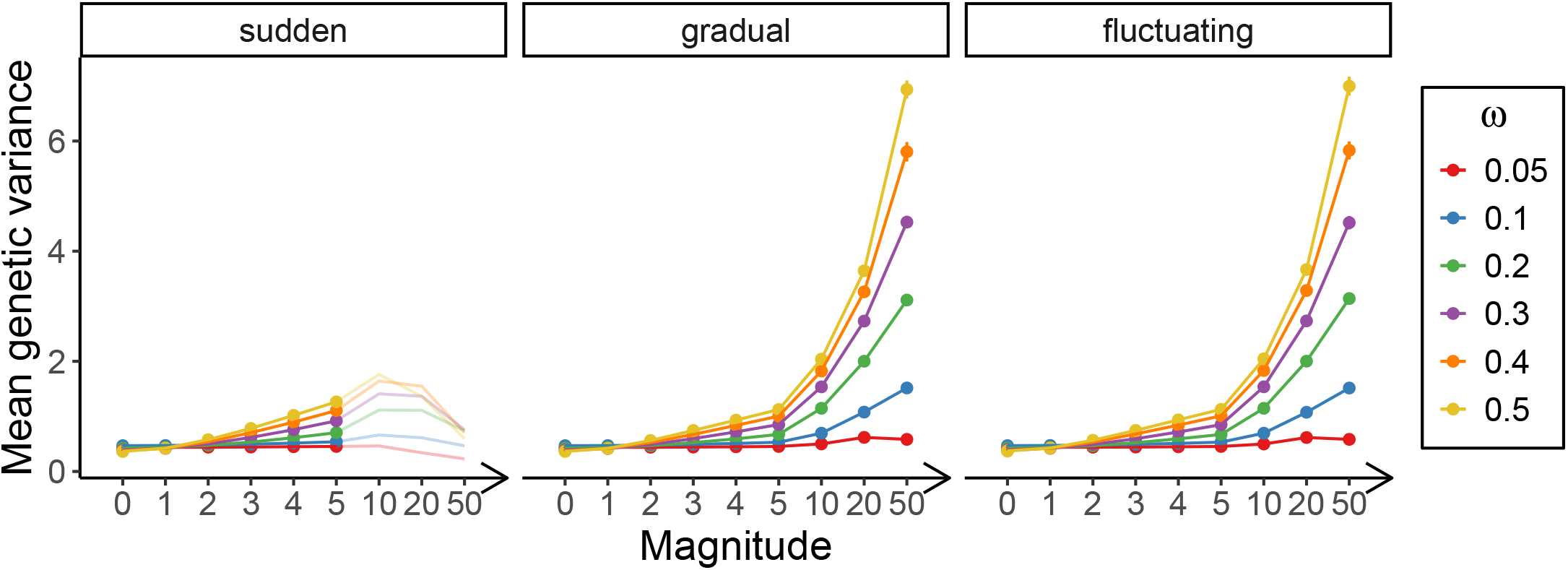
Mean genetic variance at generation 400. Each dot represents the mean genetic variance across 100 replicates, with the standard deviation indicated by error bars. Colors indicate the value of *ω*.

**Figure S8.**
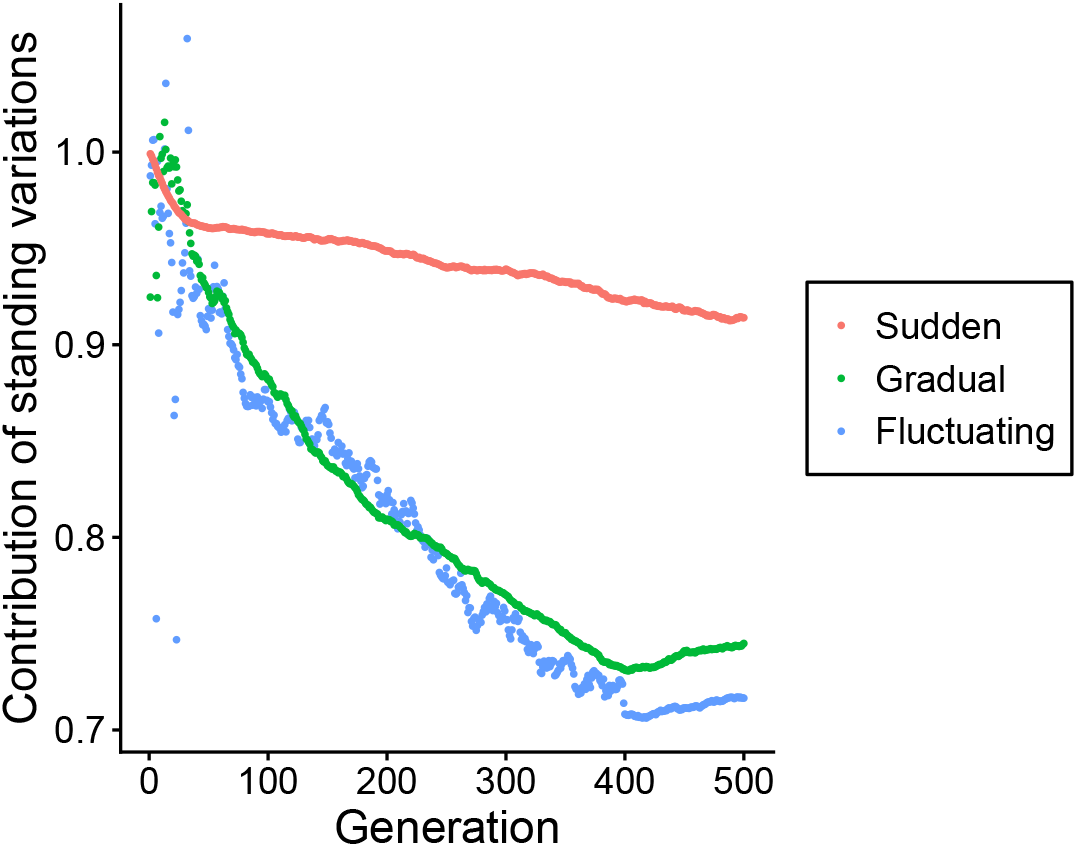
Proportion of the phenotypic contribution of standing genetic variation over time. The proportion of phenotypic changes caused by standing variants compared to the total phenotypic changes (caused by all mutations) at each generation. Colors denote shift types. Results are shown for one replicate from an example trait (magnitude=5 and *ω* = 0.3).

**Figure S9.**
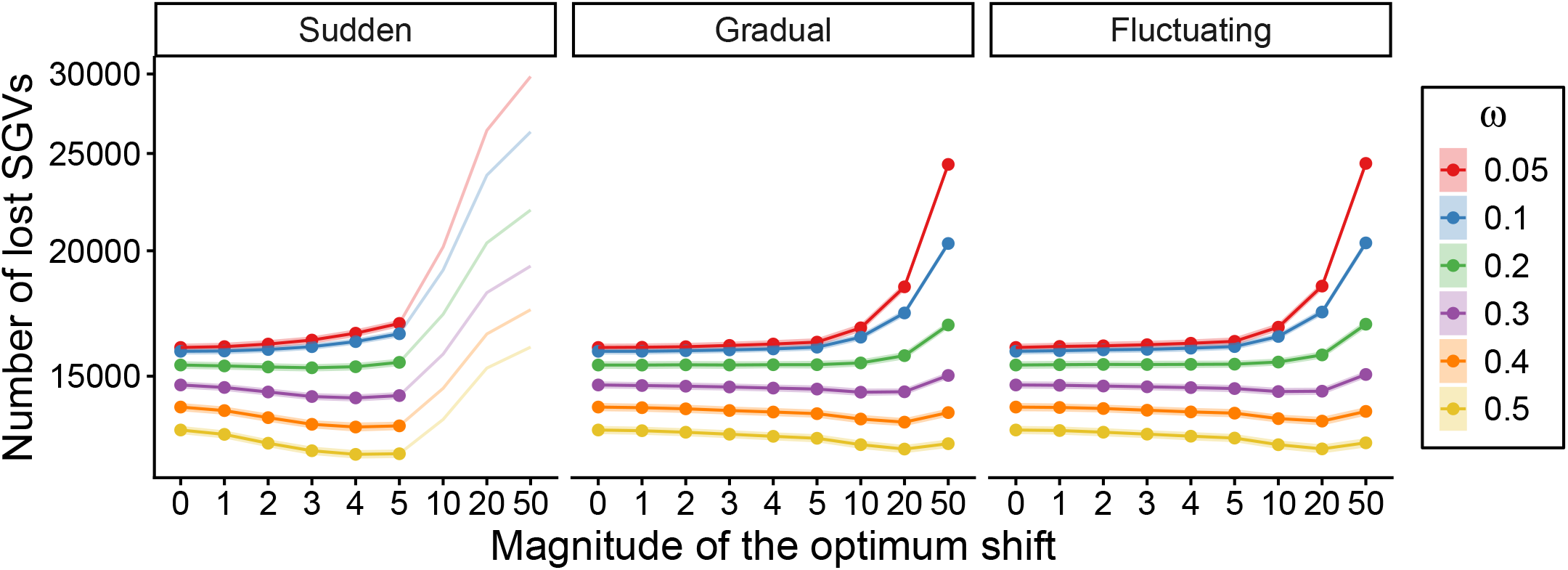
Number of lost standing genetic variants (SGVs). Each dot represents the mean number of lost standing variants across 100 replicates, with the standard deviation indicated by the shaded area. Magnitude 0 serves as a reference, representing the number of lost standing variants in static environments. The transparent lines show the results of the populations that were continuously simulated after the mean fitness fell below the extinction threshold *w <* 0.2.

**Figure S10.**
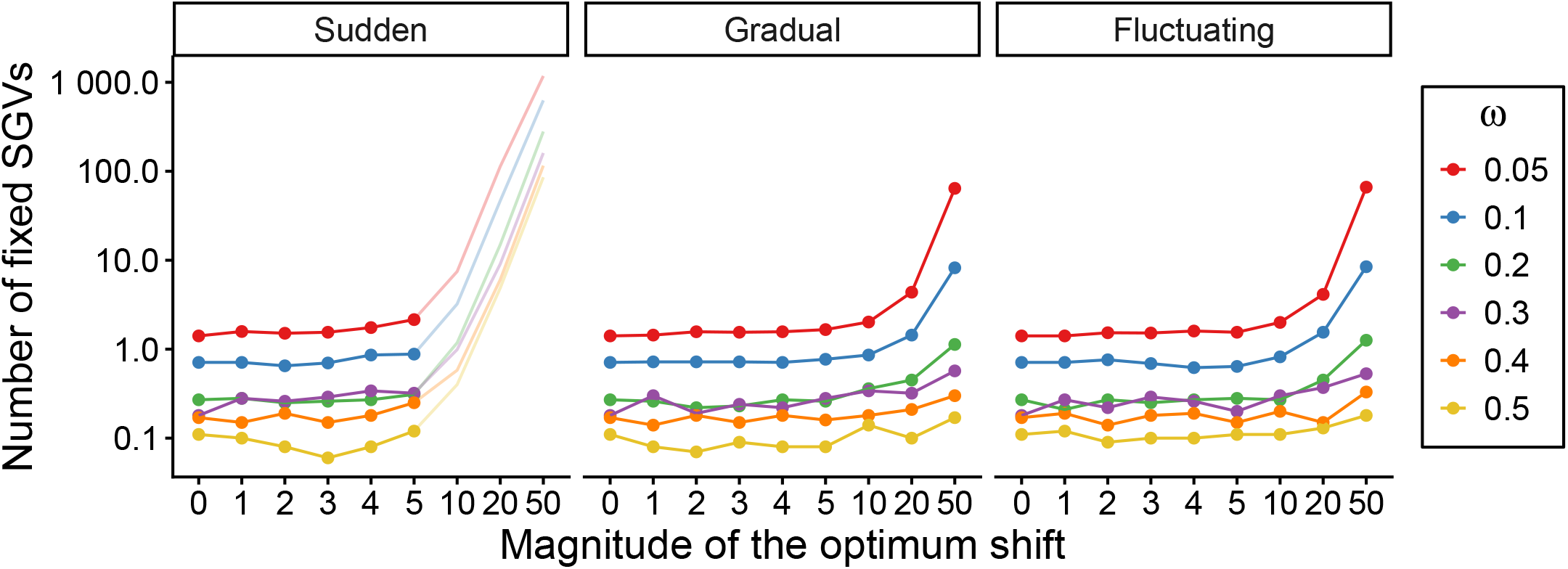
Number of fixed standing genetic variants (SGVs). Each dot represents the mean number of fixed standing variants across 100 replicates. Magnitude 0 serves as a reference, representing the mean number of fixed standing variants in static environments. The transparent lines show the results of the populations that were continuously simulated after the mean fitness fell below the extinction threshold *w <* 0.2.

**Figure S11.**
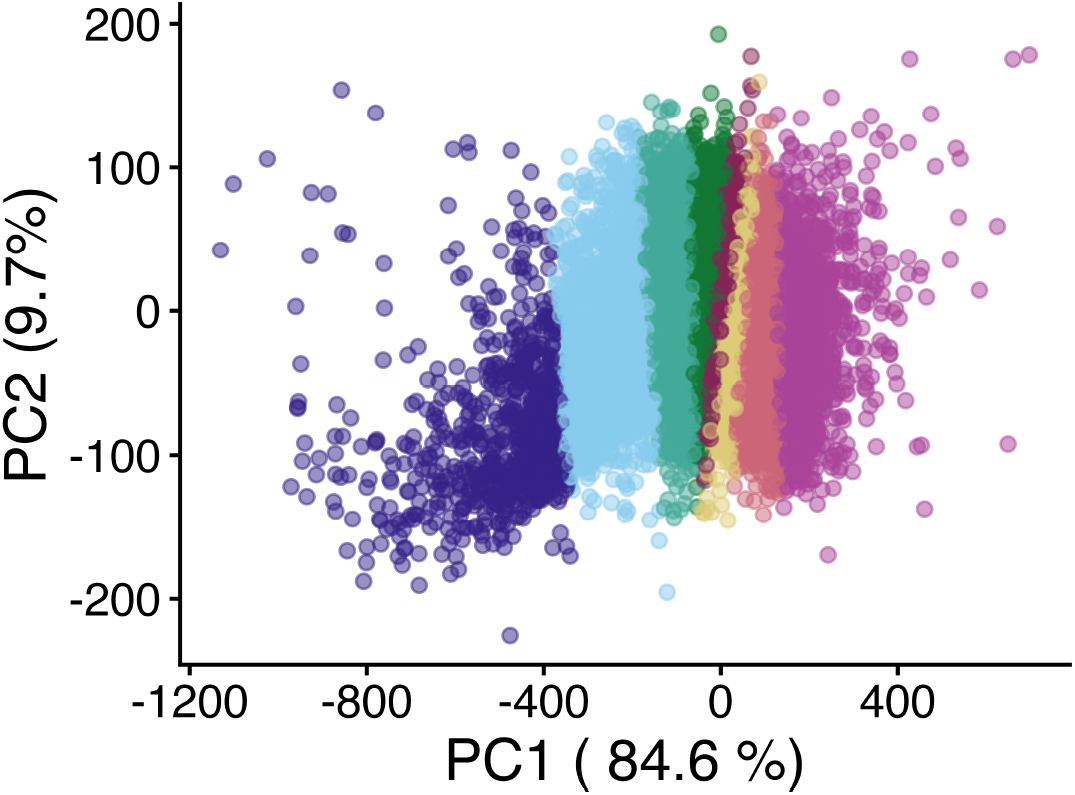
A case of PCA and k-means clustering. For populations (100 replicates) with *ω* = 0.3 under gradual environmental change with a magnitude of 50, the first two PCs of the Allele frequency trajectories are shown in the figure. Colors represent the groups of alleles based on the K-means clustering result.

**Figure S12.**
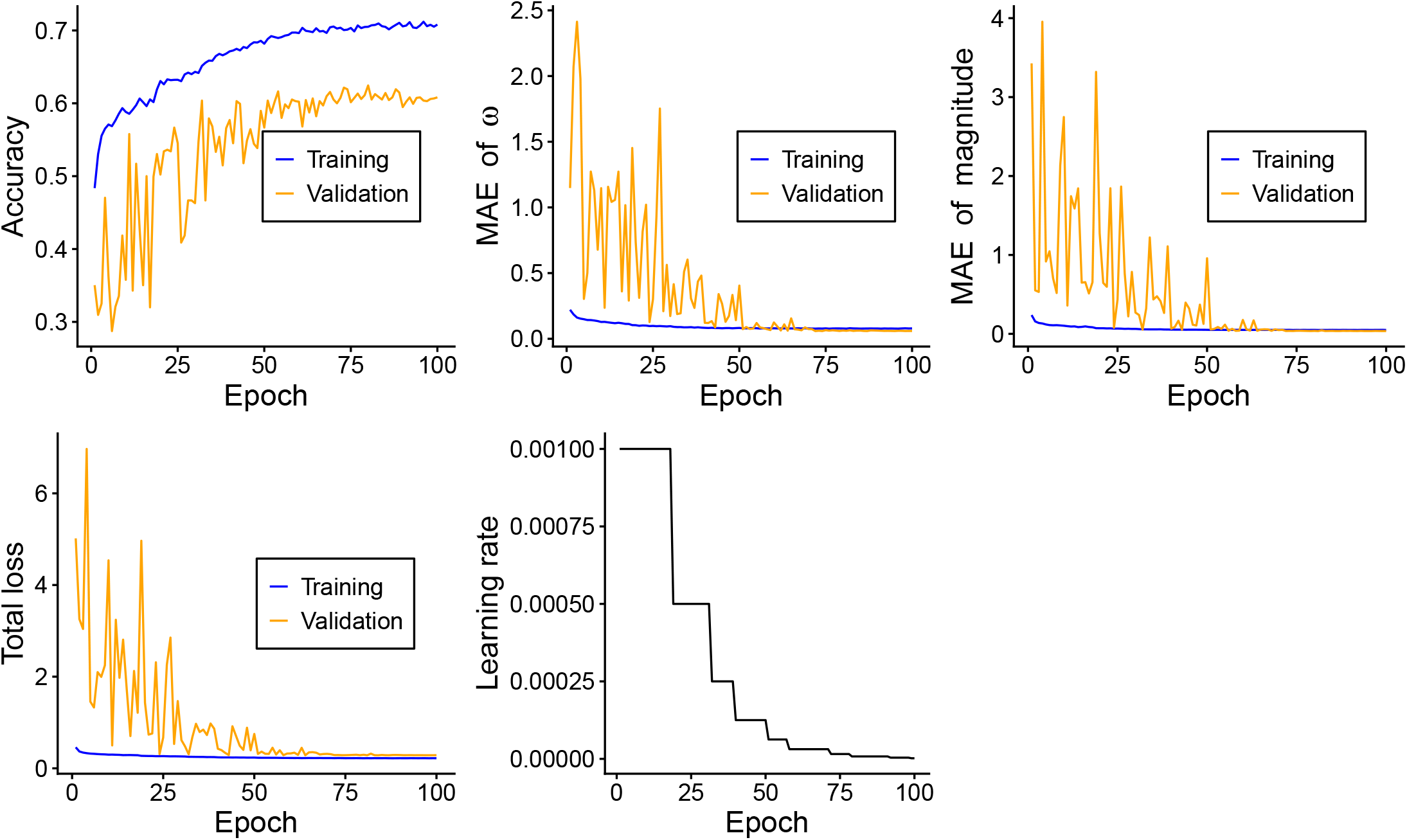
Training history for the final model with the best hyperparameter combination. The first row shows the accuracy of classifying the shift type, the MAE of the regression task for predicting the value of *ω*, and the MAE of the regression task for predicting the magnitude of environmental change, from left to right. The second row shows the total loss of all three prediction tasks and the learning rate over the course of training. The X-axis represents the course of training. Different colors represent the results on the training set and the validation set, respectively.

**Figure S13.**
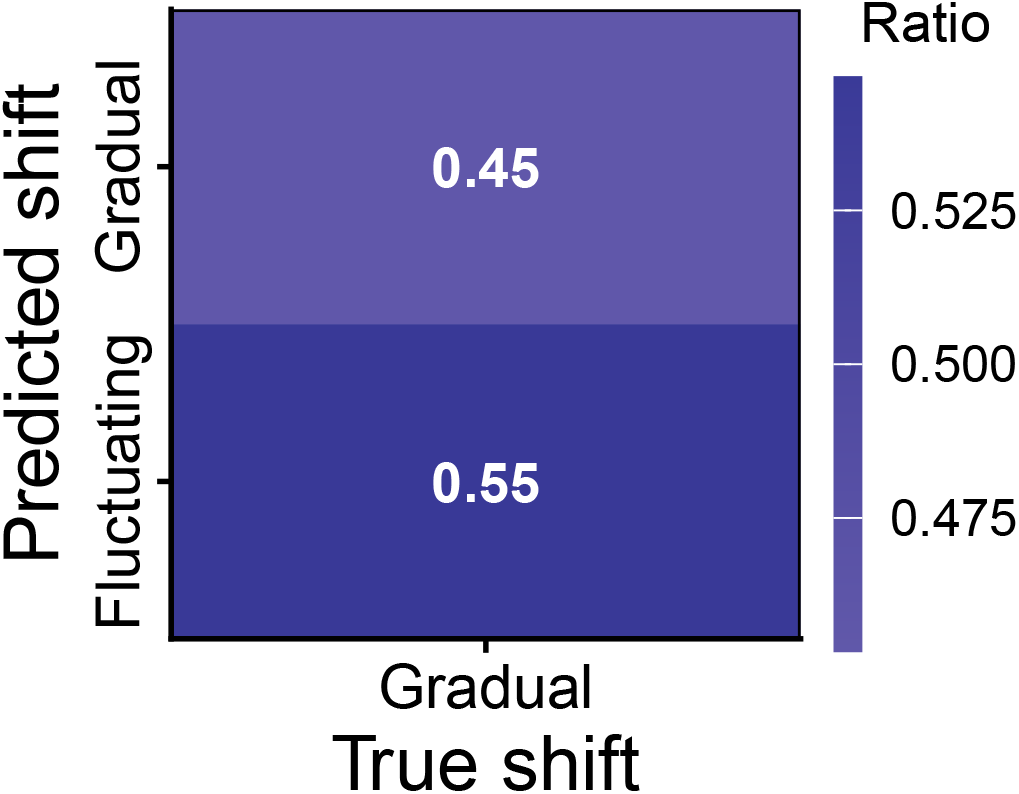
Shift type inference on the unseen dataset. The confusion matrix shows the ratio of instances in which samples from a given true shift (x-axis) were predicted as a particular shift (y-axis).

**Figure S14.**
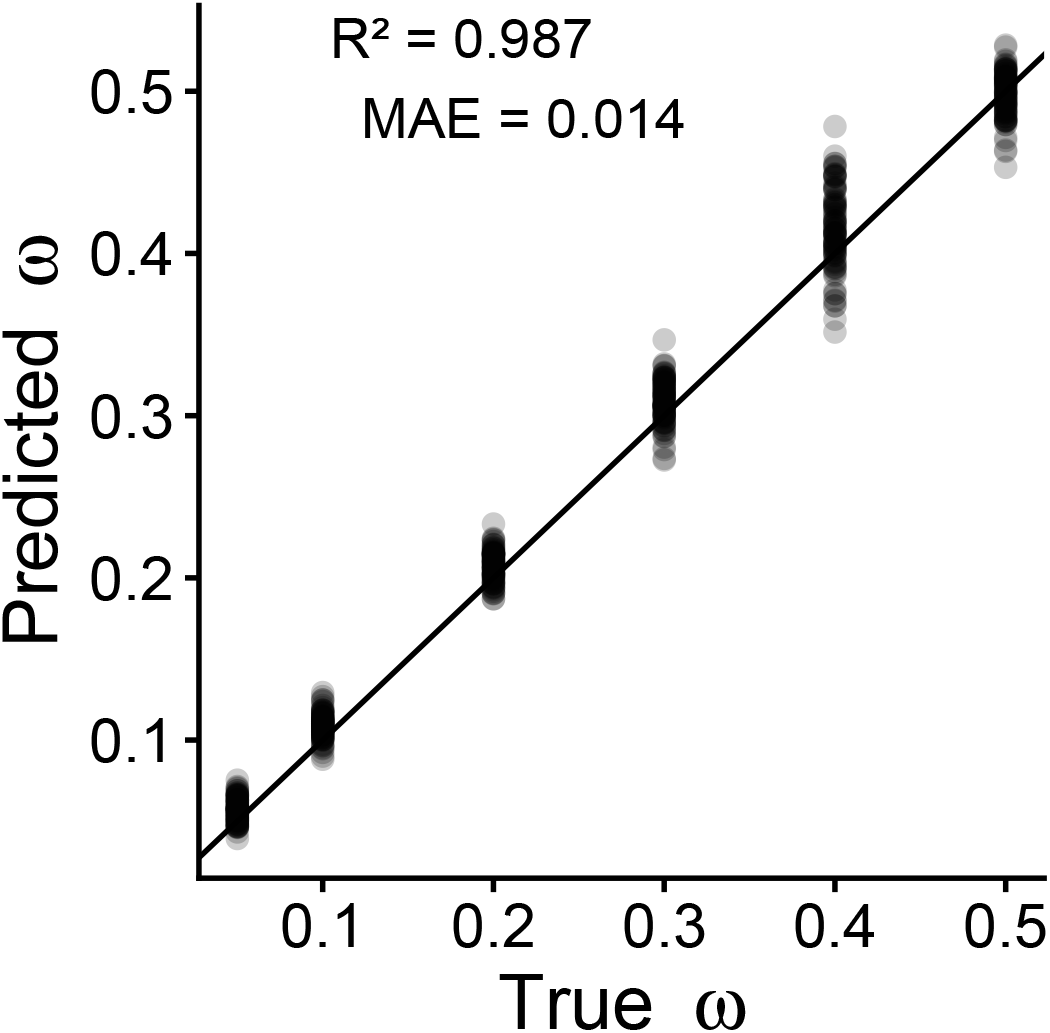
*ω* inference on the unseen dataset. Prediction results for inferring the value of *ω* of the trait on the unseen dataset. Observed and predicted values are on the x- and y-axes, respectively. The diagonal line indicates perfect agreement between predictions and true values.

**Figure S15.**
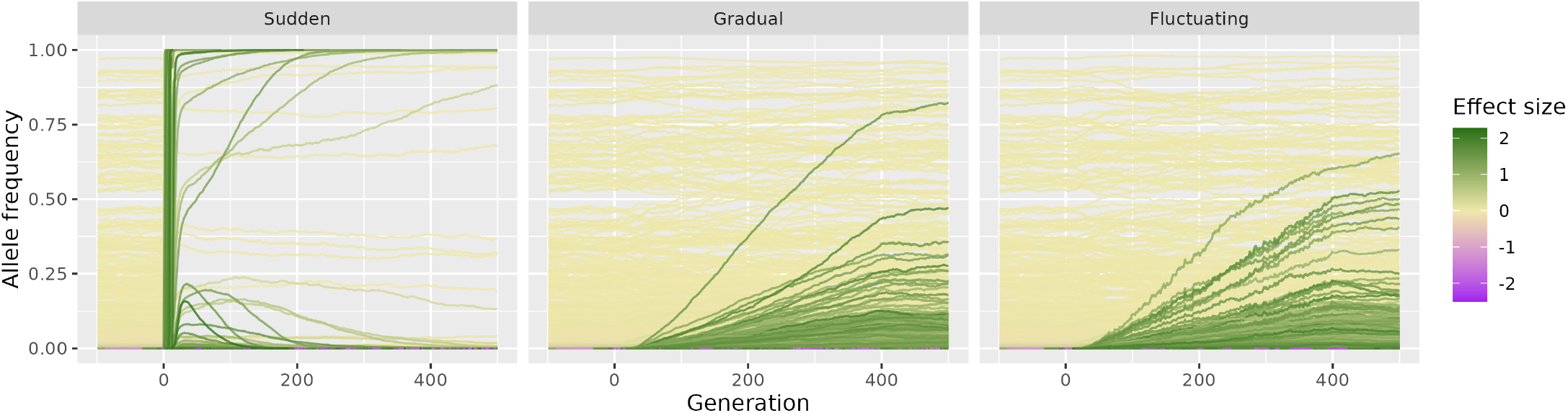
Allele frequency trajectories for traits with *ω* = 0.5 during different environmental shifts of magnitude 50. Allele frequency trajectories of all mutations during three types of environmental shifts (panels), from generation -99 to 500. Each line represents a mutation, with the color denoting the effect size of the derived allele. The result shown in each panel is from a single simulation.

## References

Abadi, M., A. Agarwal, P. Barham, E. Brevdo, Z. Chen, et al., 2015 TensorFlow: Large-Scale Machine Learning on Heterogeneous Distributed Systems.

Anderson, J. T., D. W. Inouye, A. M. McKinney, R. I. Colautti, and T. Mitchell-Olds, 2012 Phenotypic plasticity and adaptive evolution contribute to advancing flowering phenology in response to climate change. Proceedings of the Royal Society B: Biological Sciences 279: 3843–3852.

Baeckens, S. and C. M. Donihue, 2025 Evolutionary consequences of extreme climate events. Current Biology 35: R850–R864.

Barghi, N., J. Hermisson, and C. Schlötterer, 2020 Polygenic adaptation: A unifying framework to understand positive selection. Nature Reviews Genetics 21: 769–781.

Barrett, R. D. H. and D. Schluter, 2008 Adaptation from standing genetic variation. Trends in Ecology & Evolution 23: 38–44.

Battlay, P., S. Craig, A. R. Putra, K. Monro, N. P. De Silva, et al., 2025 Rapid Parallel Adaptation in Distinct Invasions of Ambrosia Artemisiifolia Is Driven by Large-Effect Structural Variants. Molecular Biology and Evolution 42: msae270.

Behrman, E. L., S. S. Watson, K. R. O’Brien, M. S. Heschel, and P. S. Schmidt, 2015 Seasonal variation in life history traits in two Drosophila species. Journal of Evolutionary Biology 28: 1691–1704.

Bergland, A. O., E. L. Behrman, K. R. O’Brien, P. S. Schmidt, and D. A. Petrov, 2014 Genomic Evidence of Rapid and Stable Adaptive Oscillations over Seasonal Time Scales in Drosophila. PLOS Genetics 10: e1004775.

Bersaglieri, T., P. C. Sabeti, N. Patterson, T. Vanderploeg, S. F. Schaffner, et al., 2004 Genetic Signatures of Strong Recent Positive Selection at the Lactase Gene. The American Journal of Human Genetics 74: 1111–1120.

Bitter, M. C., L. Kapsenberg, J.-P. Gattuso, and C. A. Pfister, 2019 Standing genetic variation fuels rapid adaptation to ocean acidification. Nature Communications 10: 5821.

Bonnet, T., M. B. Morrissey, A. Morris, S. Morris, T. H. Clutton-Brock, et al., 2019 The role of selection and evolution in changing parturition date in a red deer population. PLOS Biology 17: e3000493.

Boyle, E. A., Y. I. Li, and J. K. Pritchard, 2017 An Expanded View of Complex Traits: From Polygenic to Omnigenic. Cell 169: 1177–1186.

Buniello, A., J. A. L. MacArthur, M. Cerezo, L. W. Harris, J. Hayhurst, et al., 2019 The NHGRI-EBI GWAS Catalog of published genome-wide association studies, targeted arrays and summary statistics 2019. Nucleic Acids Research 47: D1005–D1012.

Bürger, R., 2000 The Mathematical Theory of Selection, Recombination, and Mutation. John Wiley & Sons, Chichester.

Bürger, R. and M. Lynch, 1995 Evolution and Extinction in a Changing Environment: A Quantitative-Genetic Analysis. Evolution 49: 151–163.

Carroll, S. P., H. Dingle, and S. P. Klassen, 1997 Genetic Differentiation of Fitness-Associated Traits Among Rapidly Evolving Populations of the Soapberry Bug. Evolution 51: 1182–1188.

Chen, N., I. Juric, E. J. Cosgrove, R. Bowman, J. W. Fitzpatrick, et al., 2019 Allele frequency dynamics in a pedigreed natural population. Proceedings of the National Academy of Sciences 116: 2158–2164.

Chevin, L.-M. and F. Hospital, 2008 Selective Sweep at a Quantitative Trait Locus in the Presence of Background Genetic Variation. Genetics 180: 1645.

Czorlich, Y., T. Aykanat, J. Erkinaro, P. Orell, and C. R. Primmer, 2018 Rapid sex-specific evolution of age at maturity is shaped by genetic architecture in Atlantic salmon. Nature Ecology & Evolution 2: 1800–1807.

De Vladar, H. P. and N. Barton, 2014 Stability and Response of Polygenic Traits to Stabilizing Selection and Mutation. Genetics 197: 749–767.

Durvasula, A., A. Fulgione, R. M. Gutaker, S. I. Alacakaptan, P. J. Flood, et al., 2017 African genomes illuminate the early history and transition to selfing in Arabidopsis thaliana. Proceedings of the National Academy of Sciences 114: 5213–5218.

Fairbanks, R. A. and J. Ross-Ibarra, 2025 An ancient origin of the naked grains of maize. Proceedings of the National Academy of Sciences 122: e2503748122.

Fulgione, A., C. Neto, A. F. Elfarargi, E. Tergemina, S. Ansari, et al., 2022 Parallel reduction in flowering time from de novo mutations enable evolutionary rescue in colonizing lineages. Nature Communications 13: 1461.

Futuyma, D. J., 2005 Evolution. Sinauer Associates, Sunderland (Mass.).

Grant, L., I. Vanderkelen, L. Gudmundsson, E. Fischer, S. I. Seneviratne, et al., 2025 Global emergence of unprecedented lifetime exposure to climate extremes. Nature 641: 374–379.

Grant, P. R. and B. R. Grant, 2002 Unpredictable Evolution in a 30-Year Study of Darwin’s Finches. Science 296: 707–711.

Grant, P. R. and B. R. Grant, 2006 Evolution of Character Displacement in Darwin’s Finches. Science 313: 224–226.

Greener, J. G., S. M. Kandathil, L. Moffat, and D. T. Jones, 2022 A guide to machine learning for biologists. Nature Reviews Molecular Cell Biology 23: 40–55.

Guzella, T. S., S. Dey, I. M. Chelo, A. Pino-Querido, V. F. Pereira, et al., 2018 Slower environmental change hinders adaptation from standing genetic variation. PLOS Genetics 14: e1007731.

Haller, B. C., J. Galloway, J. Kelleher, P. W. Messer, and P. L. Ralph, 2019 Tree-sequence recording in SLiM opens new horizons for forward-time simulation of whole genomes. Molecular Ecology Resources 19: 552–566.

Haller, B. C. and P. W. Messer, 2023 SLiM 4: Multispecies Eco-Evolutionary Modeling. The American Naturalist 201: E127–E139.

Hancock, A. M., S. Portalier, A. Fulgione, M. G. Stetter, and J. de Meaux, 2025 The Molecular Basis of Adaptation to Climatic Factors and Range Change in Plants. Annual Review of Ecology, Evolution, and Systematics 56: 597–621.

Hayward, L. K. and G. Sella, 2022 Polygenic adaptation after a sudden change in environment. eLife 11: e66697.

Hendry, A. P., T. J. Farrugia, and M. T. Kinnison, 2008 Human influences on rates of phenotypic change in wild animal populations. Molecular Ecology 17: 20–29.

Hendry, A. P. and M. T. Kinnison, 1999 Perspective: The Pace of Modern Life: Measuring Rates of Contemporary Microevolution. Evolution 53: 1637–1653.

Hermisson, J. and P. S. Pennings, 2005 Soft Sweeps: Molecular Population Genetics of Adaptation From Standing Genetic Variation. Genetics 169: 2335–2352.

Hoffmann, A. A. and C. M. Sgrò, 2011 Climate change and evolutionary adaptation. Nature 470: 479–485.

Höllinger, I., P. S. Pennings, and J. Hermisson, 2019 Polygenic adaptation: From sweeps to subtle frequency shifts. PLOS Genetics 15: e1008035.

Jain, K. and W. Stephan, 2015 Response of Polygenic Traits Under Stabilizing Selection and Mutation When Loci Have Unequal Effects. G3 Genes|Genomes|Genetics 5: 1065–1074.

Jain, K. and W. Stephan, 2017 Rapid Adaptation of a Polygenic Trait After a Sudden Environmental Shift. Genetics 206: 389–406.

Josephs, E. B., J. R. Stinchcombe, and S. I. Wright, 2017 What can genome-wide association studies tell us about the evolutionary forces maintaining genetic variation for quantitative traits? New Phytologist 214: 21–33.

Jost, L., 2006 Entropy and diversity. Oikos 113: 363–375.

Kelly, J. K., 2022 The genomic scale of fluctuating selection in a natural plant population. Evolution Letters 6: 506–521.

Kingma, D. P. and J. Ba, 2017 Adam: A Method for Stochastic Optimization.

Kinnison, M. T. and N. G. Hairston, 2007 Eco-Evolutionary Conservation Biology: Contemporary Evolution and the Dynamics of Persistence. Functional Ecology 21: 444–454.

Kinnison, M. T. and A. P. Hendry, 2001 The pace of modern life II: From rates of contemporary microevolution to pattern and process. Genetica 112: 145–164.

Kopp, M. and J. Hermisson, 2009a The Genetic Basis of Phenotypic Adaptation I: Fixation of Beneficial Mutations in the Moving Optimum Model. Genetics 182: 233–249.

Kopp, M. and J. Hermisson, 2009b The Genetic Basis of Phenotypic Adaptation II: The Distribution of Adaptive Substitutions in the Moving Optimum Model. Genetics 183: 1453–1476.

Kopp, M. and S. Matuszewski, 2014 Rapid evolution of quantitative traits: Theoretical perspectives. Evolutionary Applications 7: 169–191.

Lai, Y.-T., C. K. L. Yeung, K. E. Omland, E.-L. Pang, Y. Hao, et al., 2019 Standing genetic variation as the predominant source for adaptation of a songbird. Proceedings of the National Academy of Sciences 116: 2152–2157.

Lande, R., 1976 Natural Selection and Random Genetic Drift in Phenotypic Evolution. Evolution 30: 314–334.

Matuszewski, S., J. Hermisson, and M. Kopp, 2015 Catch Me if You Can: Adaptation from Standing Genetic Variation to a Moving Phenotypic Optimum. Genetics 200: 1255–1274.

Messer, P. W., 2013 SLiM: Simulating Evolution with Selection and Linkage. Genetics 194: 1037–1039.

Orr, H. A., 1998 The Population Genetics of Adaptation: The Distribution of Factors Fixed during Adaptive Evolution. Evolution 52: 935–949.

Orr, H. A., 2005 The genetic theory of adaptation: A brief history. Nature Reviews Genetics 6: 119–127.

Ossowski, S., K. Schneeberger, J. I. Lucas-Lledó, N. Warthmann, R. M. Clark, et al., 2010 The Rate and Molecular Spectrum of Spontaneous Mutations in Arabidopsis thaliana. Science 327: 92–94.

Pahujani, S., Y. Zhang, M. G. Stetter, and J. Krug, 2026 Adaptive dynamics of quantitative traits in a steadily changing environment. Genetics p. iyag168.

Pritchard, J. K. and A. Di Rienzo, 2010 Adaptation – not by sweeps alone. Nature Reviews Genetics 11: 665–667.

Pritchard, J. K., J. K. Pickrell, and G. Coop, 2010 The Genetics of Human Adaptation: Hard Sweeps, Soft Sweeps, and Polygenic Adaptation. Current Biology 20: R208–R215.

Radchuk, V., T. Reed, C. Teplitsky, M. van de Pol, A. Charmantier, et al., 2019 Adaptive responses of animals to climate change are most likely insufficient. Nature Communications 10: 3109.

Salomé, P. A., K. Bomblies, J. Fitz, R. A. E. Laitinen, N. Warthmann, et al., 2012 The recombination landscape in Arabidopsis thaliana F2 populations. Heredity 108: 447–455.

Sanderson, S., M.-O. Beausoleil, R. E. O’Dea, Z. T. Wood, C. Correa, et al., 2022 The pace of modern life, revisited. Molecular Ecology 31: 1028–1043.

Sella, G. and N. H. Barton, 2019 Thinking About the Evolution of Complex Traits in the Era of Genome-Wide Association Studies. Annual Review of Genomics and Human Genetics 20: 461–493.

Shannon, C. E., 1948 A mathematical theory of communication. The Bell System Technical Journal 27: 379–423.

Smith, J. M. and J. Haigh, 1974 The hitch-hiking effect of a favourable gene. Genetics Research 23: 23–35.

Stetter, M. G., K. Thornton, and J. Ross-Ibarra, 2018 Genetic architecture and selective sweeps after polygenic adaptation to distant trait optima. PLOS Genetics 14: e1007794.

Swarbreck, D., C. Wilks, P. Lamesch, T. Z. Berardini, M. Garcia-Hernandez, et al., 2008 The Arabidopsis Information Resource (TAIR): Gene structure and function annotation. Nucleic Acids Research 36: D1009–D1014.

Tautz, D., L. F. Pallares, L. Andersson, N. Barghi, N. Barton, et al., 2026 Beyond Mendel: A call to revisit the genotype–phenotype map through new experimental paradigms. Genetics 232: iyag024.

Tellier, A., K. Hodgins, W. Stephan, and E. Stukenbrock, 2024 Rapid evolutionary adaptation: Potential and constraints. Molecular Ecology 33: e17350.

Tuyishimire, E., M. K. Burke, and E. G. King, 2026 Dissecting fluctuating selection: A unified population and quantitative genetics framework. Genome Biology and Evolution p. evag225.

Visscher, P. M., N. R. Wray, Q. Zhang, P. Sklar, M. I. McCarthy, et al., 2017 10 Years of GWAS Discovery: Biology, Function, and Translation. Am J Hum Genet 101: 5–22.

Walther, G.-R., E. Post, P. Convey, A. Menzel, C. Parmesan, et al., 2002 Ecological responses to recent climate change. Nature 416: 389–395.

Windels, E. M., B. V. den Bergh, and J. Michiels, 2020 Bacteria under antibiotic attack: Different strategies for evolutionary adaptation. PLOS Pathogens 16: e1008431.

Wu, X., T. Bellagio, Y. Peng, L. Czech, M. Lin, et al., 2026 Rapid adaptation and extinction in synchronized outdoor evolution experiments of Arabidopsis. Science 391: eadz0777.

Yang, H., Y.-L. Li, T.-F. Xing, J.-H. Wu, T. Wang, et al., 2025 Genome-wide Parallelism Underlies Rapid Freshwater Adaptation Fueled by Standing Genetic Variation in a Wild Fish. Molecular Biology and Evolution 42.

